# Defining the RNA Modification Landscape of Multiple Myeloma Reveals METTL3-Dependent m^6^A Regulation of *NEAT1*

**DOI:** 10.64898/2026.04.04.716518

**Authors:** Prasanth Thunuguntla, Dhanusha Duraiyan, Catheryn Sizemore, Elizabeth Sulvaran-Guel, Richa Mishra, Jessica Camacho, Savannah Gonzales, Stephen Daly, Katelyn Bagwill, Dakota Colbert, Jaiyana King, Christ Samuel, Luke David-Pennington, Ashley Ki, Sydney Anderson, Carolina Bras Costa, Jin Zhang, Ravi Vij, Benjamin A. Garcia, John DiPersio, Jessica Silva-Fisher

**Affiliations:** Department of Internal Medicine, Division of Oncology, School of Medicine, Washington University in St. Louis, MO, 63110; Department of Radiation Oncology, Washington University School of Medicine, St. Louis, MO, 63108; Department of Biochemistry and Molecular Biophysics, Washington University in St. Louis, MO, 63110; Institute for Informatics, Data Science & Biostatistics (I2DB), Washington University School of Medicine, St. Louis, MO, 63110; Siteman Cancer Center, Washington University in St. Louis, MO, 63110

**Author notes:** **Correspondence** and requests for materials should be addressed to Jessica M. Silva-Fisher at. **Disclosure of Conflict of Interests/Support** R.V. (Consulting: BMS, Sanofi, Janssen, Karyopharm, Pfizer, Regenron; Research: BMS, Sanofi, Takeda), and J.D. (Consulting: Rivervest, Bluebird Bio, Vertex, HcBiosciences, SPARC; Equity-ownership WUGEN and Magenta; Research support: Macrogenics, Bioline, Incyte).

## Abstract

RNA modifications play critical roles in gene regulation. N6-methyladenosine (m^6^A) is the most abundant modification on mRNA and long noncoding RNA (lncRNA) and regulates RNA processing, stability, and translation. RNA modifications are not well characterized in multiple myeloma (MM), a plasma cell malignancy characterized by relapse and disease progression, and the contribution of m^6^A-modified lncRNAs to the disease remains unclear. Here, we define the RNA modification landscape of MM by combining mass spectrometry, Nanopore Direct RNA sequencing, and methylated RNA immunoprecipitation sequencing. We identify 20 RNA modification types and > 15,000 m^6^A sites, including sites on 2,398 lncRNAs. Among these, we validate m^6^A sites on the paraspeckle-associated lncRNA *NEAT1*. Functional studies reveal that *NEAT1* expression is regulated by the methyltransferase METTL3 and site-specific demethylation of a *NEAT1* m^6^A site reduces MM cell viability. Single-cell RNA sequencing shows consistent *NEAT1* enrichment in malignant plasma cells but minimal expression in healthy cells. These findings identify m^6^A-modified lncRNAs as key regulators of MM biology and establish *NEAT1* as an epitranscriptomically controlled driver of MM cell survival.

## BACKGROUND

RNA-based regulatory mechanisms have emerged as central determinants of gene expression control, stress adaptation, and cell-state transitions in cancer. Advances in transcriptome-wide detection technologies, including mass spectrometry,^1–5^ native molecule sequencing,^6–10^ and immunoprecipitation-based sequencing^3,11,12^ enabled the identification of more than 170 chemical modifications across coding and non-coding RNA transcripts, many of which dynamically modulate RNA fate and function.^2,7,13–19^ N^6^methyladenosine (m^6^A) is the most abundant internal RNA modification in eukaryotic mRNAs and long noncoding RNAs (lncRNAs) and regulates RNA processing, nuclear export, stability, and translation through the coordinated activity of “writers,” “erasers,” and “readers”.^20,21,22^ Dysregulation of m^6^A signaling has been linked to oncogenic gene expression programs and tumor progression across cancer types.^23–27^ Therapeutic targeting of m^6^A regulators, particularly the methyltransferase METTL3, has shown promise in preclinical and early clinical studies.^28–31^

lncRNAs comprise a major class of non-coding transcripts with diverse regulatory roles in chromatin organization, transcriptional control, and post-transcriptional gene regulation.^32–34^ Despite increasing evidence implicating lncRNAs as key drivers of cancer cell identity and disease progression,^22,34–39^ significant knowledge gaps remain regarding the contribution of RNA modifications to lncRNA function in cancer.

Here, we focus on multiple myeloma (MM), a plasma cell malignancy characterized by progressive disease and frequent relapse, reflecting the capacity of malignant cells to adapt to therapeutic pressure.^40–42^ Although current therapies for MM can induce initial responses, durable disease control remains limited, suggesting that core regulatory mechanisms governing cellular plasticity and persistence are not fully defined. While recent studies implicate METTL3-dependent m^6^A regulation in MM,^43–48^ the relevant RNA substrates and downstream targets of this regulatory pathway remain unclear. lncRNAs have also emerged as important regulators in MM, with transcriptome-wide analyses revealing widespread dysregulation associated with disease progression, proliferation, and clinical outcomes.^49–53^ Despite these data, the influence of RNA modifications on lncRNA function in MM remains unexplored.

In this study, we define the RNA modification landscape of MM and identify a network of m^6^A-modified lncRNAs. We further establish *NEAT1* as a key epitranscriptomically regulated lncRNA, uncovering a mechanism by which RNA modification controls lncRNA function to promote MM cell survival.

## METHODS

### Multiple myeloma cell lines and bone marrow patient samples

Multiple myeloma (MM) cell lines (RPMI 8226, U266B1, MM.1S, and OPM2) were generously provided by Dr. John DiPersio at Washington University in St. Louis (WashU) and cultured in RPMI 1640 media (Invitrogen, Carlsbad, CA, USA) supplemented with 15% fetal bovine serum (Invitrogen) and 1% penicillin/streptomycin (Invitrogen). MM.1R cell lines were purchased from ATCC (catalog number CRL-2975) and authenticated. All cell lines were tested for mycoplasma contamination. Relapsed/refractory MM (RRMM) patient bone marrow aspirates were obtained from the Multiple Myeloma Tissue Banking Protocol (IRB 201102270) through the Tissue Procurement Core of the Siteman Cancer Center at WashU.

For METTL3 overexpression studies, MM cell lines were transiently transfected with 2-5 µgs of METTL3 expression plasmid (Addgene, Watertown, MA, USA; Cat. #160250) or empty vector control pcDNA3.1 (Addgene, Plasmid #208051) using Lipofectamine 3000 according to the manufacturer’s protocol. Cells were incubated under standard culture conditions for 72 hours post-transfection. Total RNA was subsequently isolated using the NucleoSpin® RNA Plus Kit (Macherey-Nagel, Düren, Germany; Cat. #740984) for validation of METTL3 overexpression by RT-qPCR. Primer sequences used for gene expression analysis are listed in Supplementary Table 1.

### Detection of Mononucleoside RNA Modifications by Mass Spectrometry

Detection of global RNA modifications in MM cells was achieved using native mass spectrometry, as previously described.^2^ Total RNA was isolated from MM cell lines using the NucleoSpin® RNA Plus Kit according to the manufacturer’s instructions. Residual genomic DNA was removed using the Heat&Run gDNA Removal Kit (ArcticZymes, Tromso, Norway). RNA concentration and purity were assessed using spectrophotometric and fluorometric quantification prior to downstream processing. For global RNA modification analysis, purified RNA was processed following the nucleoside preparation workflow described in Janssen et al, 2022.^2^ Briefly, total RNA was enzymatically digested with nuclease P1 (Sigma-Aldrich, St. Louis, MO, USA) in ammonium acetate buffer at 37 °C to generate nucleoside monophosphates. Samples were subsequently treated with alkaline phosphatase (New England Biolabs, Ipswich, MA, USA) to remove phosphate groups and generate free nucleosides. The resulting nucleoside mixture was filtered and subjected to reverse-phase high-performance liquid chromatography-tandem mass spectrometry (LC-MS/MS) to quantify RNA modifications. Nucleosides were separated using a C18 analytical column under gradient elution conditions and analyzed on a high-resolution mass spectrometer operated in positive ion mode. RNA modifications were identified and quantified based on retention time and mass transitions compared with nucleoside standards.

### Nanopore Direct RNA Sequencing and Analysis

Total RNA was isolated from RPMI 8226 cell lines using the NucleoSpin® RNA Plus Kit according to the manufacturer’s instructions. Residual genomic DNA was removed using the Heat&Run gRNA Removal Kit. RNA quantity and quality were assessed prior to library preparation. Direct RNA sequencing libraries were prepared and sequenced following established protocols from the Genome Technology Access Center (GTAC) at WashU. Briefly, total RNA was used for library preparation and sequenced on an Oxford Nanopore Technologies platform according to standard direct RNA sequencing workflows. Raw Nanopore signal data (POD5 format) were basecalled using Dorado with the rna004_130bps_sup@v5.2.0 model, enabling simultaneous detection of RNA modifications via the m^6^A DRACH model. Poly(A) tail length estimation was performed during basecalling. The resulting reads were aligned to the GENCODE v44 reference transcriptome (GRCh38) using minimap2, with splice-aware parameters optimized for direct RNA sequencing (-ax splice-uf-k14), and alignments were subsequently sorted and indexed using samtools. N6-methyladenosine (m^6^A) modifications were identified using modkit, which infers RNA modification signatures from deviations in per-read ionic current signals. Putative m^6^A sites were filtered based on minimum read coverage and modification fraction thresholds, as described in the Results section. Downstream analyses were then performed to annotate m^6^A sites across transcript features and RNA biotypes.

### RNA Sequencing data processing and analysis

Publicly available single-cell RNA sequencing (scRNA-seq) count matrices from Zavidij et al.^54^ and Ledergor et al.^55^ were obtained from Gene Expression Omnibus (GEO) under accession codes GSE124310 and GSE117156, respectively. All analyses were performed in R version 4.4.2 using Seurat (version 5.2.1). Cells were filtered based on standard quality control metrics, excluding cells with less than 200 or more than 2,500 detected genes and mitochondrial transcript content greater than 5%. Counts were normalized using log normalization with a scale factor of 10,000. Normalized counts were then scaled and centered. Highly variable genes were identified using variance stabilizing transformation, with 3,000 genes retained for downstream analyses. For visualization, nonlinear dimensionality reduction was performed using Uniform Manifold Approximation and Projection (UMAP) using the uwot R package (version 0.2.2) Seurat implementation. Cell types were assigned via label transfer using the SingleR R package (version 2.8.0). Differential gene expression analysis between healthy and MM groups was performed using the Wilcoxon test implementation in Seurat and multiple testing correction was performed using Bonferroni correction. Genes with adjusted p-values lower than 0.05 were considered significant.

Bulk RNA-sequencing data from patients were obtained from the Multiple Myeloma Research Foundation (MMRF) Clinical Outcomes in Multiple Myeloma to Personal Assessment of Genetic Profiles (CoMMpass) study. Gene expression counts were used to compare newly diagnosed multiple myeloma (NDMM; n = 767) and relapsed/refractory multiple myeloma (RRMM; n = 154) samples. Differences in expression of *NEAT1* and *METTL3* were evaluated using a two-sample *t*-test. The same dataset was also used to analyze chromosomal translocations. For expression-based stratification, samples were categorized into high and low expression groups using the median expression value of each gene as the cutoff.

Methylated RNA immunoprecipitation sequencing (meRIP-Seq) data from RPMI 8226 cells with shRNA-mediated knockdown of HNRNPA2B1 compared to control shRNA treated cells was obtained from Jiang et al.^56^ lncRNA significance was determined using provided fold enrichment.

*In vivo* UV Cross-Linking Immunoprecipitation sequencing (CLIP-seq) for HNRNPA2B1 was performed in RPMI 8226 cells to identify RNA-binding sites associated with m^6^A-modified transcripts. Read quality was assessed using FastQC and MultiQC, and adapters were trimmed with Cutadapt (v5.2), removing reads <15 nt and trimming 10 bases from the 5′ end of read 1. Reads were aligned to hg38 using Bowtie2 (v2.5.4) in very-sensitive-local mode with-fr orientation. Enriched binding regions were identified using MACS2 (mfold 5–50, bandwidth 300, FDR *q* < 0.1), and crosslink sites were refined using PureCLIP (chr1-3 for training;-bc 0;-dm 8). Motif analysis was performed using RBPBench (FIMO p < 0.001; ±30 nt window). Binding profiles were visualized using IGV (v2.16). These data were used to support site-specific validation of m^6^A-associated binding on *NEAT1*.

### In vivo Cross-Linking Immunoprecipitation (CLIP) for sequencing

CLIP was performed as previously described.^50^ Briefly, cells were washed with cold PBS and crosslinked with 150 mJ/cm^2^ of UVA (254 nm) using a Stratalinker. Pellets were lysed in NP-40 buffer, incubated on ice for 10 minutes, and centrifuged to remove remaining cell debris. Supernatants were treated with 1 U/µl RNase T1 at 22°C for 30 minutes to achieve RNA fragments of 100-300 ribonucleotides. Protein G beads were pre-incubated with 5 µg isotype control antibody or anti-HNRNPA2B1 (Abcam, catalog no. ab31645, **Supplementary Table 2**) in NT2 buffer for 1 hour at room temperature. Cell lysates were added to beads and rotated for 3 hours at 4°C. Beads were washed with NP-40 buffer and treated with 20 U DNase I at 37°C for 15 minutes, then Proteinase K buffer was added to elute the proteins and crosslinked RNA fragments. RNA was extracted using phenol:chloroform:isoamyl alcohol (125:24:1) and purified using ethanol precipitation. RNA pellets were resuspended in RNase-free water and libraries were prepared for sequencing at Novogene.

### Multiplex fluorescence in situ hybridization (mFISH) and immunohistochemistry

Multiplex fluorescence in situ hybridization (mFISH) was performed using the RNAscope 2.5 HD Reagent Kit-Red assay in combination with immunohistochemistry (Advanced Cell Diagnostics [ACD], Newark, CA, USA; catalog nos. 3322360 and 323180), as previously described.^50,57^ For cytospins, bone marrow aspirates or MM cell lines were fixed onto slides, deparaffinized, treated with hydrogen peroxide, and subjected to target retrieval according to the manufacturer’s protocol. Slides were incubated overnight at 4°C with anti-Syndecan 1 (IHC138) antibody (Cell Signaling, catalog no. #30501) or an anti-m^6^A antibody (Novus Biologicals, Centennial, CO, USA; catalog no. NBP3-05657). *NEAT1*-specific RNAscope probes (ACD; catalog no. 411531) were subsequently hybridized, followed by signal amplification and Fast Red chromogenic detection. RNAscope positive control probes targeting PP1B (Catalog no. 320861) and negative control probes targeting DAPB (catalog no. 320871) were included in parallel to assess RNA integrity and background signal, respectively. Assays were considered to be acceptable if PP1B produced abundant punctate signal (> 10 dots per cell on average) with minimal background and if DAPB yielded ≤ 1 dot per 10 cells. Samples failing to meet these criteria were excluded from downstream analysis. Alexa Fluor 488-conjugated secondary antibody (Abcam, Cambridge, UK; catalog no. ab150081) was applied for one hour at room temperature in the dark. Slides were counterstained with DAPI (Sigma, St. Louis, MO, USA; catalog no. D9542), mounted using ProLong Gold Antifade Mountant (Invitrogen; catalog no. P36930), and imaged on an EVOS M5000 Imaging System (Invitrogen). RNA spot counts per cell and subcellular distribution were quantified using QuPath v0.5.1 and multiplex and compartmental analysis workflows were used to assess *NEAT1* localization across bone marrow aspirates and cell lines. Antibodies used are listed in **Supplementary Table 2**.

### Treatment of locked nucleic acid antisense oligonucleotides

Locked nucleic acid GapmeR antisense oligonucleotides (LNA ASOs) targeting *NEAT1* (*NEAT1* LNA ASO 1 or *NEAT1* LNA ASO 2) and negative control LNA ASOs were designed using the Qiagen Antisense LNA GapmeR Custom Builder (https://www.qiagen.com); sequences and catalog numbers are listed in **Supplementary Table 1**. MM cells were seeded at a density of 500,000 cells/well in six-well plates, transfected with respective LNA ASOs at 50 nM and 100 nM concentration using Lipofectamine 2000, and incubated for 72 hours. Cells were harvested, total RNA was isolated as described above, and target knockdown was validated via RT-qPCR.

### In vitro assays

Cell viability was quantified using the CellTiter-Glo® Luminescent Cell Viability Assay (Promega, Madison, WI, USA; catalog no. G7570) following the manufacturer’s protocol. Control or *NEAT1*-modified MM cell lines were seeded at 20,000 cells/well in triplicate into 96-well white opaque plates containing 50 µL of complete culture medium per well with an equal volume of CellTiter-Glo reagent added to each well. Plates were mixed by orbital shaking at room temperature for 10 minutes in the dark to stabilize the luminescent signal. Luminescence was measured using the Varioskan™ LUX microplate reader.

Cell viability and apoptosis was also measured using the ApoTox-Glo™ Triplex Assay (Promega, Madison, WI, USA; catalog no. G6320), performed according to the manufacturer’s instructions and as previously described.^50^ Control or *NEAT1*-modified MM cell lines were seeded at 20,000 cells/well in triplicate into 96-well white opaque plates containing 100 µL of complete culture medium per well. To assess cell viability and cytotoxicity, 20 µl reagent (glycyl-phenylalanyl-aminofluorocoumarin [AFC] and bis-alanylalanyl-phenylalanyl-rhodamine 110 [R110]) was added to each well, mixed for approximately 20 seconds, and incubated at 37 °C for one hour. AFC and R110, which have different excitation and emission spectra, were detected simultaneously using the Varioskan™ LUX multimode microplate reader (Thermo Fisher Scientific). To assess apoptosis, 100 µL of Caspase-Glo® 3/7 reagent (included in the ApoTox-Glo kit) was added directly to each well, mixed for approximately 20 seconds, and incubated at room temperature for one hour. Luminescence (relative luminescence units, RLU) was measured using a Varioskan™ LUX multimode microplate reader. Values were normalized to control conditions and are reported as mean ± SEM from at least three independent experiments.

### Methylated RNA immunoprecipitation (meRIP) and RT-qPCR

Total RNA was isolated as described above. Purified RNA was fragmented to an average size of ∼100 nucleotides by incubation in RNA fragmentation buffer (10 mM Tris-HCl, pH 7.0, 10 mM ZnCl₂) at 70 °C for 10 minutes, followed by immediate quenching on ice with 0.5 M EDTA. A 10% aliquot of fragmented RNA was retained as an input control. For each immunoprecipitation, 5 µg of fragmented RNA was diluted to a final volume of 500 µL in IP buffer (10 mM Tris-HCl, pH 7.4, 150 mM NaCl, 0.1% NP-40, supplemented with 0.1 U/µL RNase inhibitor). Protein A/G magnetic beads (Thermo Fisher Scientific, Waltham, MA, USA; cat. no. 88802) were incubated with 5 µg of anti-m^6^A antibody (Novus, cat. no. NBP3-05657) for one hour at 4 °C with rotation to allow antibody coupling, then washed twice with IP buffer to remove unbound antibody. Fragmented RNA was then added to antibody-conjugated beads and incubated for two hours at 4 °C with end-over-end rotation. Beads were sequentially washed once with IP buffer, twice with low-salt IP buffer, and twice with high-salt IP buffer (containing increased NaCl concentration) for 10 minutes per wash to minimize non-specific binding. Bound RNA was eluted using RLT buffer (Qiagen, Hilden, Germany). Immunoprecipitated RNA and corresponding input RNA were reverse transcribed using random hexamers and quantified by RT-qPCR with primers spanning regions of interest. Enrichment was calculated as the IP/IgG ratio following normalization to input. Each meRIP experiment included an IgG control antibody and/or a no-antibody (beads-only) control to assess background binding.

### m^6^A Dot Blot

For m^6^A dot blot analysis, RNA samples were denatured at 65 °C for five minutes then immediately placed on ice. 300 ng of denatured RNA were then spotted onto pre-wetted positively charged nylon membranes (GE Healthcare/Amersham Hybond-N+, Marlborough, MA, USA). Membranes were air-dried, UV-crosslinked at 150 mJ/cm^2^, then stained with 0.02% methylene blue in 0.3 M sodium acetate (pH 5.2) to assess RNA loading and enable normalization. After destaining with RNase-free water, membranes were blocked in 5% non-fat dry milk in TBST for one hour at room temperature and incubated overnight at 4 °C with an anti-m^6^A antibody (Sigma-Aldrich, St. Louis, MO, USA; cat. no. ABE572). Membranes were washed with TBST then incubated with an HRP-conjugated anti-rabbit secondary antibody (Cell Signaling Technology, Danvers, MA, USA; cat. no. 7074S) for one hour at room temperature. Membranes were treated with enhanced chemiluminescence (ECL) reagents and signals were detected using standard chemiluminescent detection systems. m^6^A signal intensity was quantified and normalized to methylene blue staining to control for RNA loading. All experiments were performed in biological triplicate.

### Targeted m^6^A demethylation of NEAT1

Site-specific m^6^A demethylation of *NEAT1* was performed using a CRISPR-dCas13a-based RNA editing system. A catalytically inactive *Leptotrichia wadei* Cas13a (dCas13a) was fused to the m^6^A demethylase FTO (dCas13a–FTO), as previously described,^58^ and was generously provided by Dr. Andrew Hutchins. CRISPR RNAs (crRNAs) targeting *NEAT1* were designed and cloned under a U6 promoter by VectorBuilder (Chicago, IL, USA). A non-targeting (NT) crRNA was used as a negative control. Lentiviral particles were produced in HEK293T cells (ATCC, Manassas, VA, USA) by co-transfection of 7.5 µg of dCas13a–FTO or control lentiviral construct together with packaging plasmids: 2 µg psPAX2 (Addgene plasmid #12260) and 1 µg pMD2.G (VSV-G; Addgene plasmid #12259), using standard lipid-based transfection reagents. Viral supernatants were harvested 72 hours post-transfection, filtered through a 0.45 µm membrane, and used to transduce target MM cell lines in the presence of 8 µg/mL polybrene (Sigma-Aldrich, St. Louis, MO, USA; cat. no. TR-1003-G). Stable cell populations were selected using puromycin (0.5 µg/mL) for five days. Successful site-specific demethylation was verified by using meRIP RT-qPCR to confirm reduced m^6^A occupancy at the targeted *NEAT1* site without alteration of total *NEAT1* transcript levels.

### METTL3 knockdown and inhibitor treatment

MM cell lines were seeded at a density of 4-5 × 10^5^ cells per well in six-well plates. For genetic depletion, cells were transfected with two small interfering RNAs (siRNAs) targeting *METTL3* (Invitrogen, Thermo Fisher Scientific, Waltham, MA, USA; Cat. #4392420, Assay ID: s32141) using Lipofectamine 3000 (Invitrogen, Thermo Fisher Scientific) according to the manufacturer’s protocol. Cells were incubated for 72 hours following transfection prior to downstream analyses. siRNA sequences are listed in **Supplementary Table 1**.

For pharmacologic inhibition, cells were treated with the METTL3 inhibitor STM2457 (MilliporeSigma, Burlington, MA, USA; Cat. #SML3360) at final concentrations of 0, 10, or 20 µM for 72 hours. Control cells were treated with an equivalent volume of DMSO vehicle. All treatments were performed under standard cell culture conditions. Following treatment, cells were harvested and processed for downstream analyses, including the ApoTox-Glo™ Triplex Assay (Promega), CellTiter-Glo® Luminescent Cell Viability Assay (Promega), and m^6^A dot blot analysis.

## Data availability statement

All sequencing data is available at GEO under accession numbers GSE326610 and GSE326744 for Nanopore direct RNA sequencing and CLIP-seq respectively. In accordance with the MMRF data use agreement, the authors are prohibited from distributing raw data, derivatives, or results obtained from the MMRF CoMMpass study on secondary or third-party databases. Instead, these data are publicly available to qualified researchers for non-commercial scientific research through the MMRF Virtual Lab™ (VLab) platform. Access can be requested by registering directly on the VLab portal https://themmrf.org/for-researchers/data-access-via-virtual-lab/. All other data are available within this article and its Supplementary Information files. All other data supporting the findings of this study are available from the corresponding author upon reasonable request.

## Ethics approval and consent to participate

All methods were performed in accordance with relevant guidelines and regulations. This study was conducted under the supervision of the Washington University Human Rights Protections Office and in compliance with all relevant guidelines and regulations for clinical research. All participants in CoMMpass (NCT01454297) provided informed consent for that study. As these analyses used only de-identified patient-level data, they were determined to be exempt from human subjects research requirements. RRMM patient bone marrow aspirates were obtained from participants following informed consent and registration on our internal Multiple Myeloma Tissue Banking Protocol (protocol number 201102270).

## RESULTS

### Global RNA modification profiling in multiple myeloma

Given growing evidence that RNA modifications play critical roles in cancer,^27^ we assessed the epitranscriptomic landscape of MM by profiling global RNA modifications in five MM cell lines (RPMI 8226, U266B1, OPM2, MM.1S, and MM.1R) that reflect the biological diversity observed in MM. Global RNA modifications were quantified using native mass spectrometry-based nucleoside analysis (**Figure 1a**).^2^ Many RNA modifications, for instance adenosine derivatives (e.g., m^6^A, m^1^A, and m^6^Am), are structurally and mass-similar, often exhibiting isobaric properties that necessitate simultaneous detection for accurate discrimination. We detected 20 distinct RNA modifications across the five MM cell lines, including six adenosine, five cytidine, two guanosine, and seven uridine modifications (**Figure 1b** and **Supplementary Table 3**).

**Figure 1:**
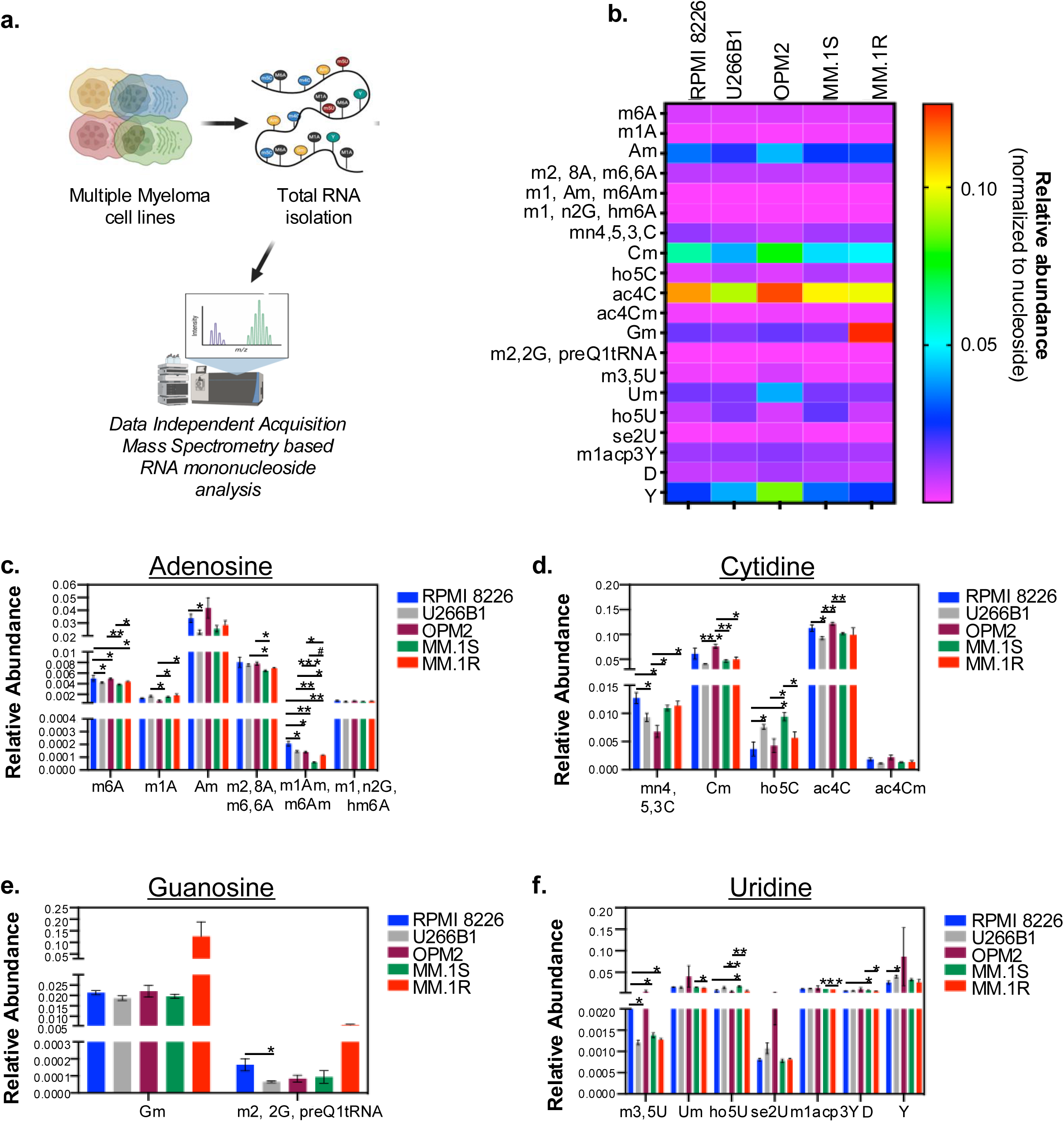
Global RNA modification profiling using mass spectrometry in myeloma cell lines. **a.** Process of isolating total RNA from myeloma cell lines for native mass spectrometry. **b**. Heatmap showing abundance of RNA modifications identified in five MM cell lines. **c–f.** Relative abundance of specific **c.** adenosine, **d.** cytosine, **e.** guanosine, and **f.** uridine RNA modifications across MM cell lines. *p value < 0.05, **p value < 0.005, ***p value < 0.0005

Comparative analysis revealed that several modifications were present at similar levels across all MM cell lines, including m^1^,n2G, hm^6^A, ac^4^Cm, Gm, and se^2^U. In contrast, there were significant differences in levels of the adenosine modification m^1^Am, m^6^Am across nearly all pairwise comparisons (RPMI 8226 versus U266B1 p = 0.03; RPMI 8226 versus MM.1S p = 0.001; RPMI 8226 versus OPM2 p = 0.02; RPMI 8226 versus MM.1R p = 0.007; U2266B1 versus MM.1S p = 0.0003; U226B1 versus MM.1R p = 0.01; MM.1S versus OPM2 p = 0.0005; MM.1S versus MM.1R p = 7.51 x 10^-5^; and OPM2 versus MM.1R p = 0.03), with only U266B1 versus OPM2 showing no differences (**Figure 1c**). Notably, dexamethasone-sensitive MM.1S and dexamethasone-resistant MM.1R cells exhibited the most pronounced differences across the modification landscape, with m¹Am, m^6^Am representing the top differentially modified nucleosides between these two cell lines. Additional significant differences between MM.1S and MM.1R cells were observed for D (p = 0.001), m¹acp³Y (p = 0.0009), ho⁵U (p = 0.003), and m^6^A (p = 0.02; **Figures 1c-f**).

m^6^A is the most abundant internal modification in eukaryotic mRNAs and lncRNAs. While m^6^A levels were comparable between RPMI 8226 and OPM2, U266B1 and MM.1S, and OPM2 and MM.1R, significant differences were detected in several comparisons, including RPMI 8226 versus U266B1 (p = 0.04), RPMI 8226 versus MM.1S (p = 0.01), RPMI 8226 versus MM.1R (p = 0.04), MM.1S versus OPM2 (p = 0.008), and MM.1S versus MM.1R (p = 0.02; **Figure 1c**).

Together, these data reveal extensive epitranscriptomic heterogeneity across MM cell lines and identify distinct RNA modification signatures associated with dexamethasone sensitive and resistant MM cell lines, supporting a potential role for RNA modifications in disease progression in MM.

### Direct RNA sequencing identifies specific RNA targets with m^6^A RNA modifications

Following the identification of diverse RNA modifications in MM cells (**Figure 1**), we focused on transcripts harboring m^6^A, the predominant internal RNA modification enriched on lncRNAs and a key regulator of RNA fate and oncogenic gene expression programs. To accomplish this, we performed Nanopore direct RNA sequencing (direct RNA-seq) in RPMI 8226 cells to detect transcriptome-wide m^6^A sites (**Figure 2a**). To ensure high confidence m^6^A site detection, we applied a >50% modification threshold using ModKit, which calculates the ratio of modified to unmodified reads at each position. At these identified sites, we observed an average depth of 9.86 modified reads per site compared to 3.07 unmodified reads (**Figure 2b**). Protein-coding (mRNA) transcripts exhibited the highest coverage, with 11.78 reads for m^6^A sites and 3.70 reads for unmodified adenines, whereas lncRNAs showed lower coverage overall, with 3.25 reads for m^6^A sites and 0.65 reads for unmodified adenines (**Figure 2b**).

**Figure 2:**
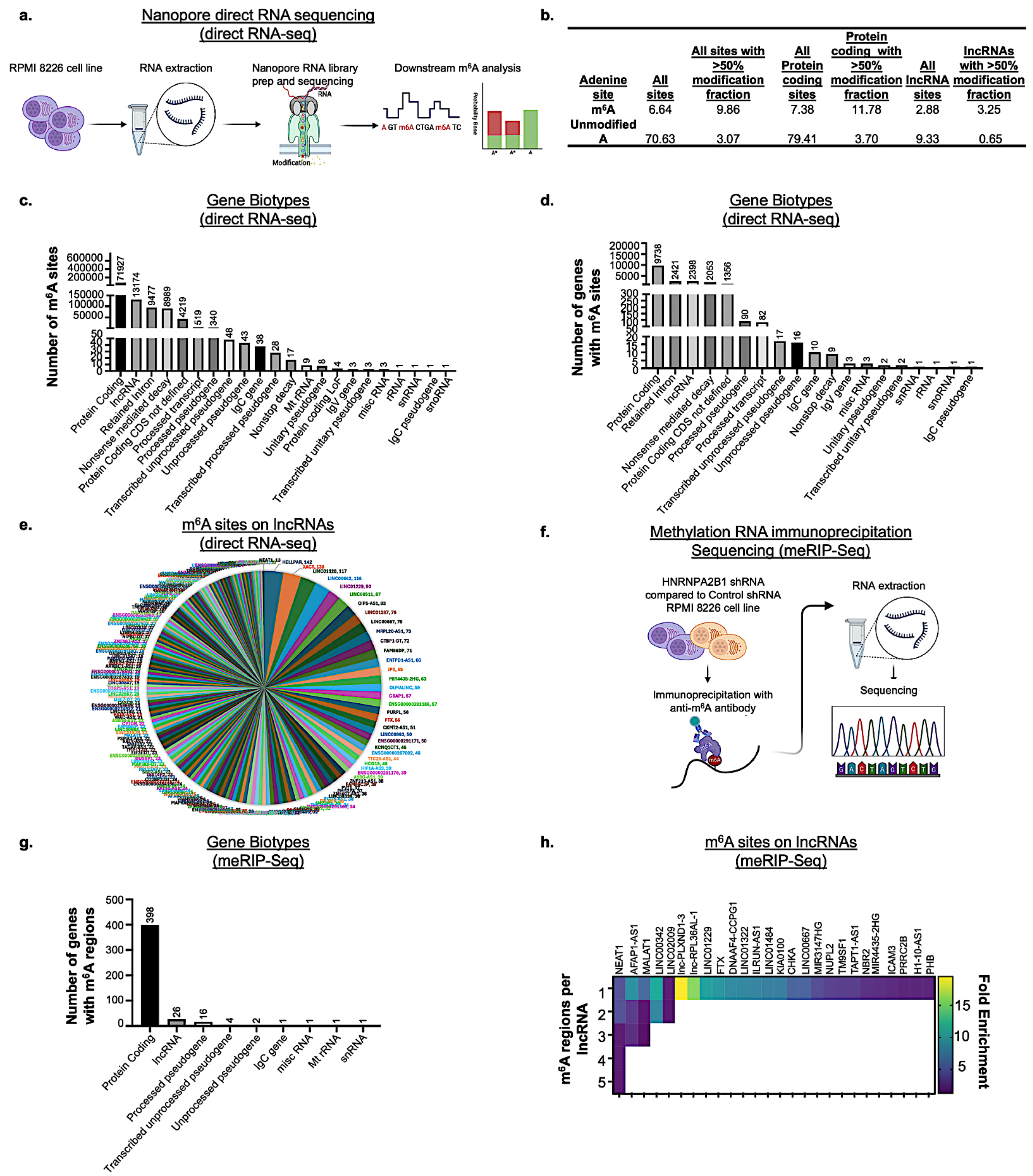
Global RNA modification profiling using sequencing for site-specific modifications in myeloma cell lines. **a.** Process of Nanopore Direct RNA Sequencing (direct RNA-seq). **b.** m^6^A modification fractions detected by direct RNA-seq for all adenine sites, protein coding sites, and lncRNA sites. **c.** Total number of m^6^A sites detected by direct RNA-seq per gene biotype. **d.** Total number of genes with m^6^A sites with > 50% modification fraction detected by direct RNA-seq per gene biotype. **e.** Top m^6^A-modified lncRNAs detected by direct RNA-seq. **f**. Process of methylation RNA immunoprecipitation (meRIP-Seq) comparison of HNRNPA2B1 shRNA treated vs control shRNA treated RPMI 8226 cells. **g.** RNA subtypes identified from methylation RNA immunoprecipitation sequencing (meRIP-Seq) in control shRNA treated RPMI 8226 cells when comparing to HNRNPA2B1 shRNA treated cell lines. **h.** Fold enrichment of lncRNAs identified as having potential m^6^A modifications and numbers of m^6^A regions identified from meRIP-Seq.

Transcriptome-wide analysis revealed that the majority of m^6^A sites (66%, n = 71,927) were located within protein-coding genes, followed by lncRNAs (12%, n = 13,174) and retained introns (9%, n = 9,477; **Figure 2c**). When categorized by gene, 53% (n = 9,738) of m^6^A-modified transcripts corresponded to protein-coding genes, followed by retained introns (13%, n = 2,421) and lncRNAs (13%, n = 2,398; **Figure 2d**). Additional m^6^A sites were detected in transcripts annotated as nonsense-mediated decay targets, processed transcripts, pseudogene variants, immunoglobulin constant (IgC) and variable (IgV) genes, and a subset of small noncoding RNAs (**Figure 2d**).

Protein-coding transcripts containing m^6^A sites included previously reported oncogenic targets, such as *MYC*^26,59,60^ and *ILF3*.^56,61^ Interestingly, the top transcript with the highest coverage and modified fractions identified was *G3BP2*, a protein previously reported to selectively exclude or “repel” m^6^A-modified RNAs (**Supplementary Table 4**).^62^ Of the 2,398 lncRNAs identified with m^6^A sites, the majority (> 690 lncRNAs; 29%) contained a single m^6^A site, while the remaining transcripts contained two to six sites. The highest number of m^6^A sites (n = 142) was detected in the lncRNA *Hepatocellular Carcinoma-Upregulated Long Noncoding RNA Promoting Autophagy and Regeneration (HELLPAR*; **Figure 2e and Supplementary Table 5**). Notably, several lncRNAs previously implicated in cancer and MM biology were also identified as harboring multiple m^6^A sites, including *Nuclear Enriched Abundant Transcript 1* (*NEAT1)*, *Metastasis Associated Lung Adenocarcinoma Transcript 1 (MALAT1)*, and *Plasmacytoma Variant Translocation 1 (PVT1*; **Figure 2e and Supplementary Table 5**).^46,63^ Overall, these results characterize a landscape of m^6^A modifications in a MM cell line and show that while most modifications occur on protein-coding transcripts, a substantial fraction is present on lncRNAs.

### lncRNA-specific m^6^A modifications in multiple myeloma

Several well-characterized lncRNAs, including *MALAT1*, *TUG1*, and *NEAT1*, are known to contain multiple m^6^A modifications in solid tumors, which regulate lncRNA function and promote metastatic progression.^64–66^ Notably, m^6^A modification of *NEAT1* has been shown to promote prostate, bone, and lung cancer metastasis in mouse models through regulation of Ser2 phosphorylation of RNA polymerase II.^67^ These observations suggest that m^6^A-dependent regulation of lncRNAs may represent a conserved mechanism contributing to tumor progression.

Given our identification of widespread RNA modifications in MM (**Figure 1**), including lncRNA-associated modifications (**Figure 2b-e**), we sought to validate specific lncRNA targets harboring m^6^A modifications in the context of MM using methylated RNA immunoprecipitation sequencing (meRIP-seq) data, an antibody-based enrichment approach that can quantify m^6^A modifications directly. HNRNPA2B1 protein is a well-studied m^6^A reader that is expressed in MM.^56,68^ We analyzed a publicly available meRIP-seq dataset comparing RPMI 8226 cells with shRNA-mediated knockdown of the m^6^A reader HNRNPA2B1 and control shRNA-treated cells (**Figure 2f**).^69^ HNRNPA2B1 is expressed in MM and has been shown to mediate m^6^A-dependent RNA regulation.^56,68^ This analysis revealed that the majority of identified m^6^A sites localized to protein-coding transcripts (88%, n = 398), with additional enrichment observed on lncRNAs (6%, n = 26), processed pseudogenes (4%, n = 16), unprocessed pseudogenes (1%, n = 6), and less than 1% on snRNAs, miscRNAs, mitochondrial RNAs, and immunoglobulin constant region transcripts (**Figure 2g**).

*NEAT1*, previously reported to contain m^6^A sites in cancer,^20,66,67,70–89^ exhibited the highest number of m^6^A-modified regions detected by meRIP-Seq among these lncRNAs with five distinct peaks (region 1-5 fold enrichment = 4.57, 4.52, 1.89, 1.88, and 1.70; **Figure 2h**). Other lncRNAs with prominent m^6^A enrichment included AFAP1 antisense RNA 1 (*AFAP1-AS1*), which contained three m^6^A-modified regions (fold enrichment = 8.99, 3.32, and 2.70), and *MALAT1*, which also contained three m^6^A-modified regions (fold enrichment = 6.17, 1.65, and 1.25; **Figure 2h**). The single most highly enriched m^6^A region identified in RPMI 8226 cells mapped to *lnc-PLXND1-3* (fold enrichment = 19.50; **Figure 2h**).

Despite the lower site-level resolution of this antibody-based enrichment approach, this analysis complements direct RNA-seq by enabling orthogonal validation of m^6^A-modified transcripts.

### *NEAT1* is a highly m^6^A-modified lncRNA in multiple myeloma

To establish the functional relevance of m^6^A-modified lncRNAs, we sought to validate candidate transcripts by integrating orthogonal datasets. We compared m^6^A modified lncRNAs identified by direct RNA-seq (**Figure 2e**) and meRIP-seq (**Figure 2g–h**), revealing 19 overlapping regions with > 50% modification fractions (**Figure 3a, Supplementary Table 6**). Direct RNA-seq identified a greater number of m^6^A sites overall, consistent with its higher resolution and sensitivity to low-stoichiometry modifications relative to meRIP-seq, which identifies broader regions of methylation enrichment. *NEAT1* contained the highest number of m^6^A sites validated by meRIP-seq of all overlapping candidate lncRNAs. In addition, although m^6^A modifications on *NEAT1* are reported in other cancer contexts, its m^6^A modification status and functional relevance in MM remain largely unexplored.

**Figure 3:**
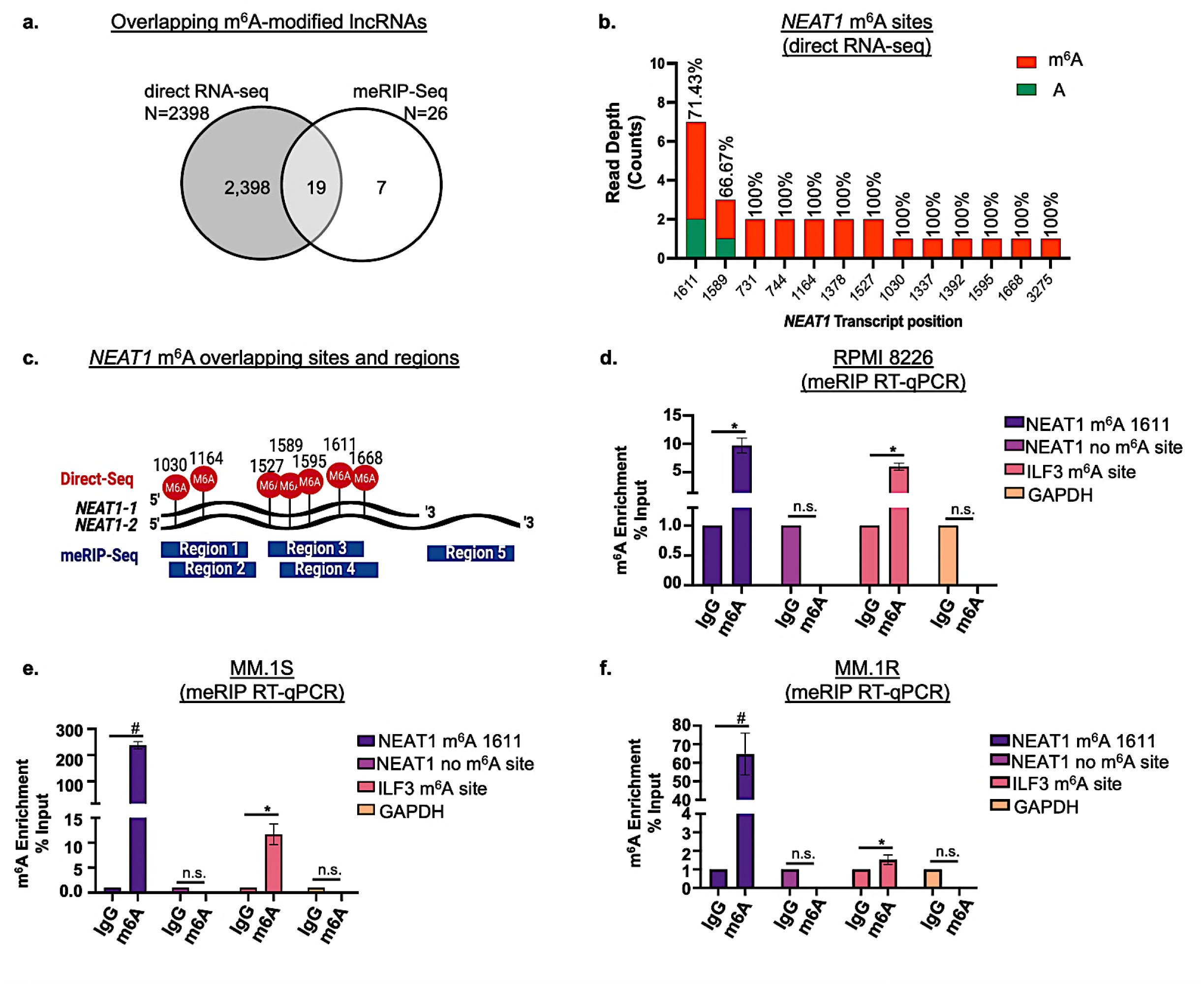
m^6^A sites on *NEAT1* lncRNA in myeloma. **a.** Venn diagram showing overlap of m^6^A-modified lncRNAs identified by direct RNA-seq and meRIP-Seq. **b.** Read depth of *NEAT1* m^6^A sites identified by direct RNA-seq. **c.** *NEAT1* m^6^A site positions identified by both direct RNA-seq (red) and meRIP-Seq (blue). **d-f.** meRIP RT-qPCR data validating m^6^A site 1611 on *NEAT1* in **d.** RPMI 8226, **e.** MM.1S, and **f.** MM.1R relative to negative control IgG. *ILF3* mRNA serves as positive control and *GAPDH* serves as negative control. *fold enrichment > 5, ^#^fold enrichment > 50, n.s. not significant.

Reflecting this, we performed site-level analysis of *NEAT1* using direct RNA-seq. This analysis identified 13 m^6^A sites on *NEAT1*, including several highly modified positions (site 1611, 71.43% m^6^A modified; site 1589, 66.67% m^6^A modified; and 11 sites with 100% modification; **Figure 3b**), with site-level concordance between platforms shown in **Figure 3c**. We used the sequence-based m^6^A site predictor SRAMP^90^ to further prioritize functionally relevant sites on *NEAT1*, which identified site 1611 as a very high-confidence m^6^A consensus site (score = 0.803; **Supplementary Figure 1**).

We focused on *NEAT1* site 1611 for validation due to its high m^6^A modification fraction in direct RNA-seq data and high-confidence computational prediction as an m^6^A consensus site (**Supplementary Figure 2)**. We conducted UV crosslinking immunoprecipitation sequencing (CLIP-seq) and CLIP RT-qPCR in wild type RPMI 8226 cells and confirmed binding of the m^6^A reader HNRNPA2B1 at this site (**Supplementary Figure 3a-b**). meRIP RT-qPCR across multiple MM cell lines (RPMI 8226, MM.1S, and MM.1R) demonstrated consistent robust enrichment at *NEAT1* m^6^A site 1611 (mean fold enrichment: RPMI 8226 = 9.7; MM.1S = 237.3; MM.1R = 64.72) and at positive control *ILF3* m^6^A sites (mean fold enrichment RPMI 8226 = 5.96; MM.1S = 11.68; MM.1R = 1.52), whereas no enrichment was observed in a non-methylated *NEAT1* region or negative control GAPDH (**Figures 3d–f**).

Together, these data show that *NEAT1* is a highly m^6^A-modified lncRNA in MM and identify site 1611 as a relevant epitranscriptomic regulatory site.

### Site-specific removal of m^6^A from *NEAT1* suppresses multiple myeloma cell growth

To investigate the functional role of m^6^A modifications on *NEAT1* in MM directly, we used a catalytically inactive RNA-targeting CRISPR effector (dCas13a) fused to the m^6^A demethylase FTO to enable site-specific removal of m^6^A modifications without altering the underlying nucleotide sequence.^58^ We used the dCas13a-FTO complex guided by short CRISPR RNAs (crRNAs) targeting *NEAT1* site 1611 to assess the functional consequences of loss of m^6^A modifications at this specific site (**Figure 4a**). Removal of m^6^A modifications at site 1611 (dCas13a-FTO/NEAT1 m^6^A-1611) did not significantly alter total *NEAT1* transcript abundance compared with non-targeting controls (dCas13a-FTO/crNT) in either RPMI 8226 or MM.1S cells (**Figure 4b and c**). In addition, we treated the dCas13a-FTO/NEAT1 m^6^A-1611 cells with the RNA polymerase inhibitor Actinomycin D and did not detect a significant change in stability (**Supplementary Figure 4**), indicating that site-specific demethylation of site 1611 does not impact *NEAT1* stability or transcription. meRIP RT-qPCR analysis showed m^6^A enrichment at site 1611 was significantly lower in dCas13a-FTO/NEAT1 m^6^A-1611 cells, as compared to dCas13a-FTO/crNT controls, in both RPMI 8226 (fold change > 40; **Figure 4d**) and MM.1S cells (fold change > 5; **Figure 4e**).

**Figure 4:**
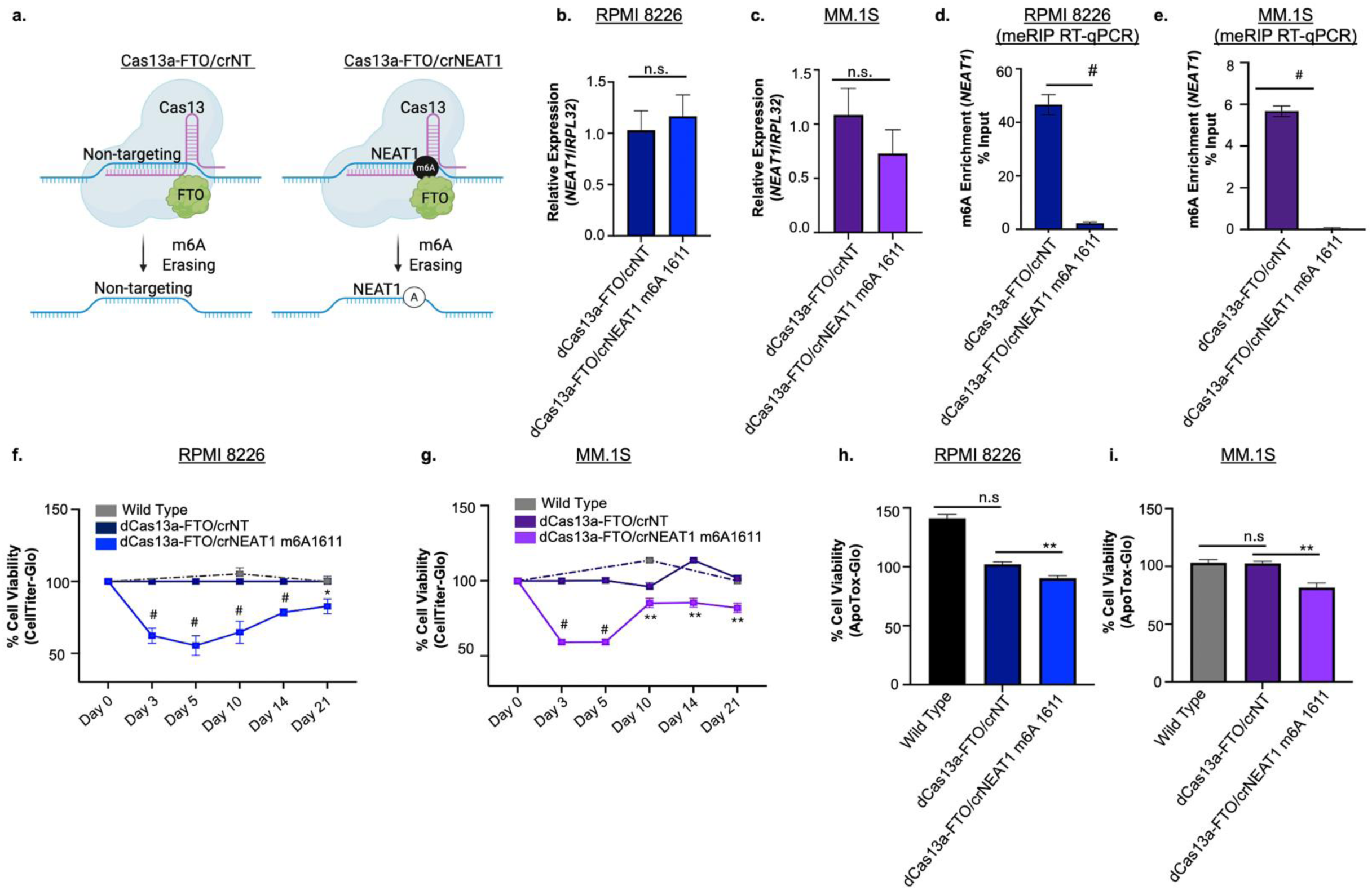
Demethylating m^6^A site 1611 on *NEAT1* decreases viability. **a.** CRISPR-dCas13a-based system to demethylate m^6^A sites using a catalytically dead LwaCas13a (dCas13a) fused to FTO demethylase domain (dCas13a-FTO) guided by NEAT1 crRNA (crNEAT1) or non-targeting crRNA (crNT) as control. **b and c.** Expression of *NEAT1* in dCas13a-FTO/crNT and dCas13a-FTO/crNEAT1 m^6^A 1611 treated **b.** RPMI 8226 and **c.** MM.1S cells. **d and e**. Methylation RNA immunoprecipitation (meRIP) RT-qPCR showing decreased enrichment of m^6^A on site 1611 in dCas13a-FTO/crNEAT1 m^6^A 1611 treated **d.** RPMI 8226 and **e.** MM.1S cells, as compared to dCas13a-FTO/crNT treated cells. **f and g.** Long-term viability assay showing a decrease in viability in dCas13a-FTO/crNEAT1 m^6^A 1611 treated **f.** RPMI 8226 cells and **g.** MM.1S cells, as compared to wild type and dCas13a-FTO/crNT treated cells. **h and i.** Decreased viability in dCas13a-FTO/crNEAT1 m^6^A 1611 treated **h.** RPMI 8226 cells. **i.** MM.1S cells, as compared to dCas13a-FTO/crNT treated cells, measured by ApoTox-Glo assay. *p value < 0.05, **p value < 0.005, ***p value < 0.0005, n.s. not significant.

Functional analysis showed that site-specific removal of m^6^A from *NEAT1* site 1611 significantly reduced MM cell viability. RPMI 8226 dCas13a-FTO/NEAT1 m^6^A-1611 cells exhibited a robust reduction in viability compared with both wild-type and dCas13a-FTO/crNT controls, as measured by CellTiter-Glo (Day 3 p = 4.22 × 10⁻⁵; Day 5 p = 5.43 × 10⁻⁵; Day 10 p = 0.0001; Day 14 p = 1.25 × 10⁻⁵; Day 21 p = 0.0007; **Figure 4f**) and ApoTox-Glo assays (p = 0.001; **Figure 4h**). A comparable decrease in viability was observed in MM.1S dCas13a-FTO/NEAT1 m^6^A-1611 cells (Day 3 p = 0.0004; Day 5 p = 4.31 × 10⁻⁶; Day 10 p = 9.46 × 10⁻⁶; Day 14 p = 0.002; Day 21 p = 0.002; **Figure 4g**), and ApoTox-Glo assays (p = 0.008; **Figure 4i**).

Together, these findings demonstrate that m^6^A modification at a single, defined site on *NEAT1* is required to sustain MM cell viability, establishing a direct, site-specific functional role for m^6^A-modified *NEAT1* in promoting MM progression.

### METTL3 promotes myeloma cell survival and regulates *NEAT1*

METTL3, the catalytic core of the m^6^A methyltransferase complex, is upregulated in MM and promotes malignant phenotypes through m^6^A-dependent regulation of oncogenic mRNAs and non-coding RNAs, including pathways controlling proliferation, apoptosis, and stem-like properties.^46–48^ METTL3-mediated m^6^A deposition modulates RNA stability, processing, and translation in MM and other cancers, including on lncRNAs that regulate chromatin organization, transcriptional output, and cellular stress adaptation.^43–45^ Prior studies have reported METTL3-dependent m^6^A modification of *NEAT1*,^44,66,70,83,91–95^ supporting a potential regulatory interaction. Based on these observations, we next assessed the role of METTL3 in regulating *NEAT1* m^6^A modification in MM.

Analysis of bulk RNA sequencing data from newly diagnosed MM (NDMM, n = 767) and relapsed/refractory MM (RRMM, n = 154) patient samples from the MMRF CoMMpass study revealed that *METTL3* expression is significantly higher in RRMM compared with NDMM patient samples (p = 6.1 × 10⁻⁵; **Supplementary Figure 5a**). We next assessed *METTL3* expression in plasma cells and B cells isolated from CD138⁻ bone marrow fractions in single-cell RNA sequencing (scRNA-seq) data from a study by Zavidij et al.,^54^ which included nine healthy control samples and seven MM patient samples (**Figure 5a**), and an independent scRNA-seq dataset in plasma cells and B cells isolated from CD138⁻ bone marrow fractions from Ledergor et al.,^55^ which included 11 healthy donor samples and 12 symptomatic MM patient samples. This analysis showed no significant differences in *METTL3* expression between healthy and MM samples (**Figure 5a and b**; **Supplementary Figure 5b and c**). Further assessment of RRMM patient samples with genetic translocations revealed overall higher number of patients containing low expression of *METTL3* associated with Amp1q and higher expression of *METTL3* in patients with t(11;14) cases (**Supplementary Figure 5d**). This suggests that *METTL3* upregulation may reflect disease progression, cell state-specific regulation, or underlying genetic alterations that are not detectable at the single-cell transcriptomic level, potentially compounded by the limited number of patients analyzed.

**Figure 5:**
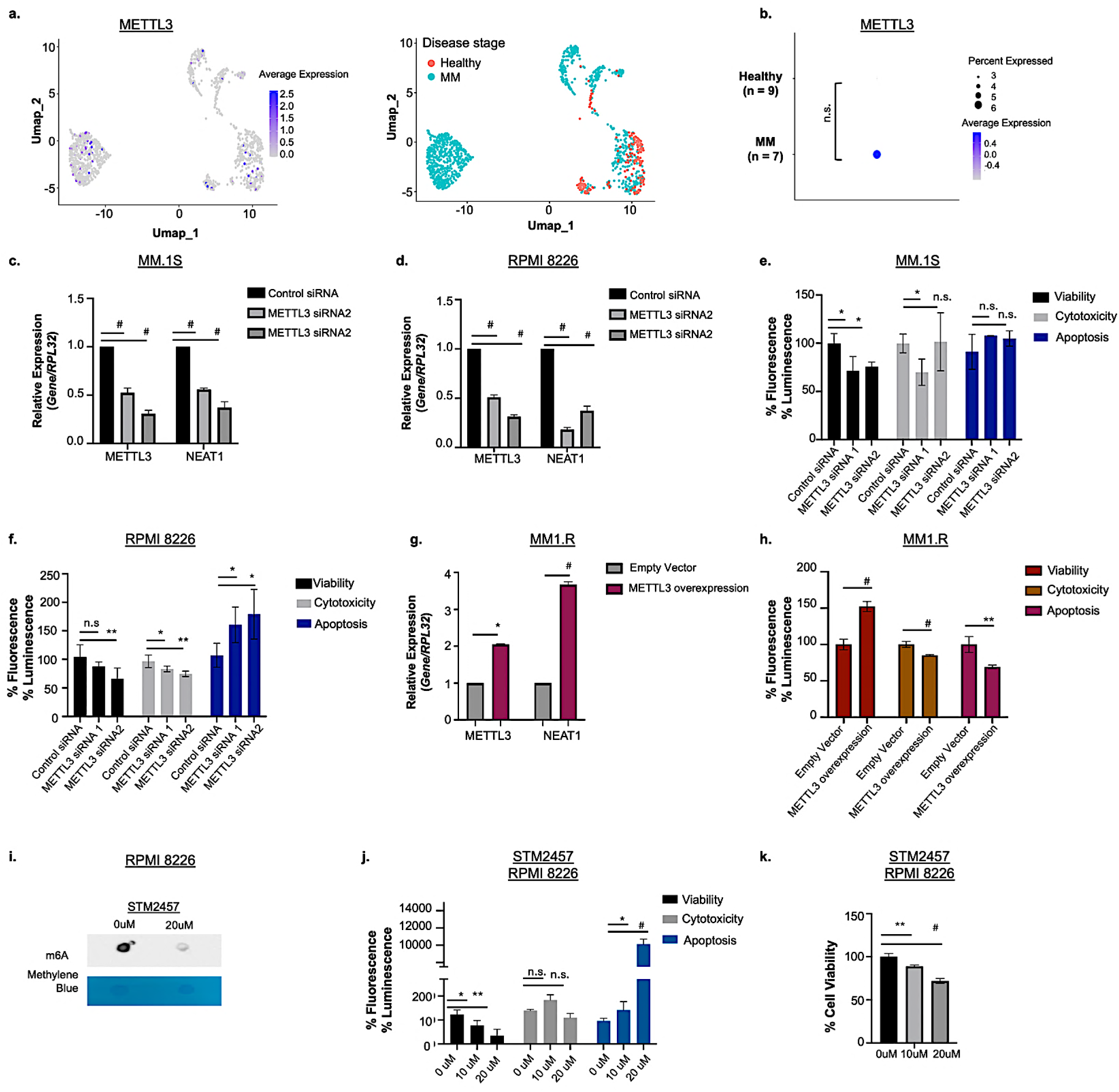
M*E*TTL3 is expressed in MM and regulates *NEAT1* expression in myeloma cells. **a.** UMAP plot of B and plasma single-cell RNA sequencing data for *METTL3* (left plot) from healthy (red) and multiple myeloma (MM) patients (blue) (right plot). **b.** Expression of *METTL3* in MM samples compared to healthy samples. **c and d**. Expression of *METTL3* and *NEAT1* following small interfering RNA (siRNA)-mediated *METTL3* knockdown in **c.** MM.1S cells and **d.** RPMI 8226 cells. **e.** Decreased viability in MM.1S cells following siRNA-mediated *METTL3* knockdown, as measured by ApoTox-Glo assay. **f.** Decreased viability and increased apoptosis in RPMI 8226 cells following siRNA-mediated *METTL3* knockdown, as measured by ApoTox-Glo assay. **g.** Expression of *METTL3* and *NEAT1* in MM.1R cells with a *METTL3* overexpression vector. **h.** Increased viability and decreased apoptosis in MM.1R cells with a *METTL3* overexpression vector, as measured by ApoTox-Glo assay. **i.** m^6^A blot showing global m^6^A levels in MM cells after 72-hour treatment with METTL3 inhibitor STM2457. **j.** Decreased viability and increased apoptosis in RPMI 8226 cells with increasing concentrations of STM2457 treatment, as measured by ApoTox-Glo assay **k.** Decreased viability in RPMI 8226 cells with increasing concentrations of STM2457 treatment, as measured by Celltiter-Glo Assay. *p value < 0.05, **p value < 0.005, ***p value < 0.0005, #p value < 0.00005, n.s. not significant

Prior studies demonstrated that METTL3 catalyzes m^6^A modification of *NEAT1*, thereby enhancing its stability and oncogenic function across multiple cancer contexts.^66,67,70,78,83^ To directly test this relationship in MM, we treated MM cell lines with two independent small interfering RNAs (siRNAs) targeting *METTL3* and assessed relative expression of both *METTL3* and *NEAT1*. siRNA-mediated *METTL3* knockdown significantly decreased *NEAT1* expression in both MM.1S cells that contain high endogenous expression of *METTL3* (*METTL3*: siRNA1 p = 0.0006, siRNA2 p = 4.6 × 10⁻⁵; *NEAT1*: siRNA1 p = 9.76 × 10⁻⁶, siRNA2 p = 0.0004) and RPMI 8226 cells (*METTL3*: siRNA1 p = 0.0001, siRNA2 p = 4.3 × 10⁻⁶; *NEAT1*: siRNA1 p = 1.53 × 10⁻⁵, siRNA2 p = 0.0001; **Figure 5c and d**). Endogenous *METTL3* expression levels in MM cell lines are shown in **Supplementary Figure 5e**. Functional analysis showed that siRNA-mediated *METTL3* knockdown significantly reduced cell viability in MM.1S cells relative to non-targeting control siRNA (siRNA1 p = 0.02; siRNA2 p = 0.01; **Figure 5e**), with more modest effects in RPMI 8226 cells (siRNA2 p = 0.008; **Figure 5f**). Increased apoptosis was observed in RPMI 8226 cells following *METTL3* knockdown (siRNA1 p = 0.04; siRNA2 p = 0.03; **Figure 5f**), with significant reduction in cytotoxicity. We also assessed the effects of *METTL3* overexpression in MM.1R cells that contained low endogenous expression of *METTL3,* (**Figure 5g**) and showed that it significantly increased cell viability (p = 0.000046) and decreased both cytotoxicity (p = 0.0003) and apoptosis (p = 0.0014; **Figure 5h**).

To complement our knockdown and overexpression analyses, we pharmacologically inhibited METTL3 using STM2457, a selective METTL3 catalytic inhibitor and predecessor of STC-15.^96^ STM2457 treatment in RPMI 8226 cells reduced global m^6^A levels (**Figure 5i**), decreased cell viability in a dose-dependent manner (10 μM p = 0.02, 20 μM p = 0.001; **Figure 5j**) and increased apoptosis (10 μM p = 0.029; 20 μM p = 2.72 e^-11^; **Figure 5j**). A decrease in cell viability was also detected using a CellTiter-Glo assay (10 μM p = 0.008, 20 μM p = 0.0004; **Figure 5k**). Together, these data establish METTL3 as a functional regulator of *NEAT1* expression and MM cell survival and demonstrate that both genetic and pharmacologic inhibition of METTL3 disrupts m^6^A-dependent oncogenic RNA programs in MM.

To determine whether METTL3 directly mediates m^6^A modifications on *NEAT1* in MM specifically, we knocked down *METTL3* expression using siRNA in U266B1 cells and performed meRIP RT-qPCR to quantify site-specific m^6^A enrichment. We found that siRNA-mediated *METTL3* knockdown induced a 7.09-fold decrease in m^6^A enrichment at *NEAT1* site 1611 and an 8.28-fold decrease at the positive control *ILF3* m^6^A site, relative to non-targeting control siRNA (**Supplementary Figure 6**). These findings support the functional relevance of METTL3-mediated methylation and are consistent with observations that treatment of MM cells with the METTL3 inhibitor STM2457 reduced global m^6^A levels and decreased cell viability.

Together, these data demonstrate that METTL3 is required for m^6^A deposition on *NEAT1*. Our findings that both genetic knockdown and pharmacologic inhibition of METTL3 impair MM cell survival highlight its key role as an epitranscriptomic regulator of oncogenic lncRNA function in MM.

### *NEAT1* expression and relevance in multiple myeloma

*NEAT1* is broadly expressed across normal tissues, with particularly high expression in epithelial and glandular organs (**Supplementary Figure 7**). Dysregulated *NEAT1* expression has been implicated in the pathogenesis of multiple cancers, including MM, and has well established oncogenic roles associated with promoting poor prognosis, therapeutic resistance, and disease progression.^54,95,97–115^ Despite this, *NEAT1* expression dynamics in MM have not been previously defined at a single-cell level, particularly in the context of m^6^A modification.

To address this, we assessed *NEAT1* expression at single-cell resolution by analyzing scRNA-seq data from Zavidij et al.^54^ We found significant upregulation of *NEAT1* in MM plasma and B cells compared to healthy controls (log₂FC = 0.64, FDR = 6.7 × 10⁻⁴³; **Figures 6a and b**). Consistent with this observation, analysis of the scRNA-seq dataset from Ledergor et al.^55^ showed a statistically significant, but more modest increase in *NEAT1* expression in MM (log₂FC = 0.20, FDR = 2.5 × 10⁻³⁶; **Supplementary Figure 8a**).

**Figure 6:**
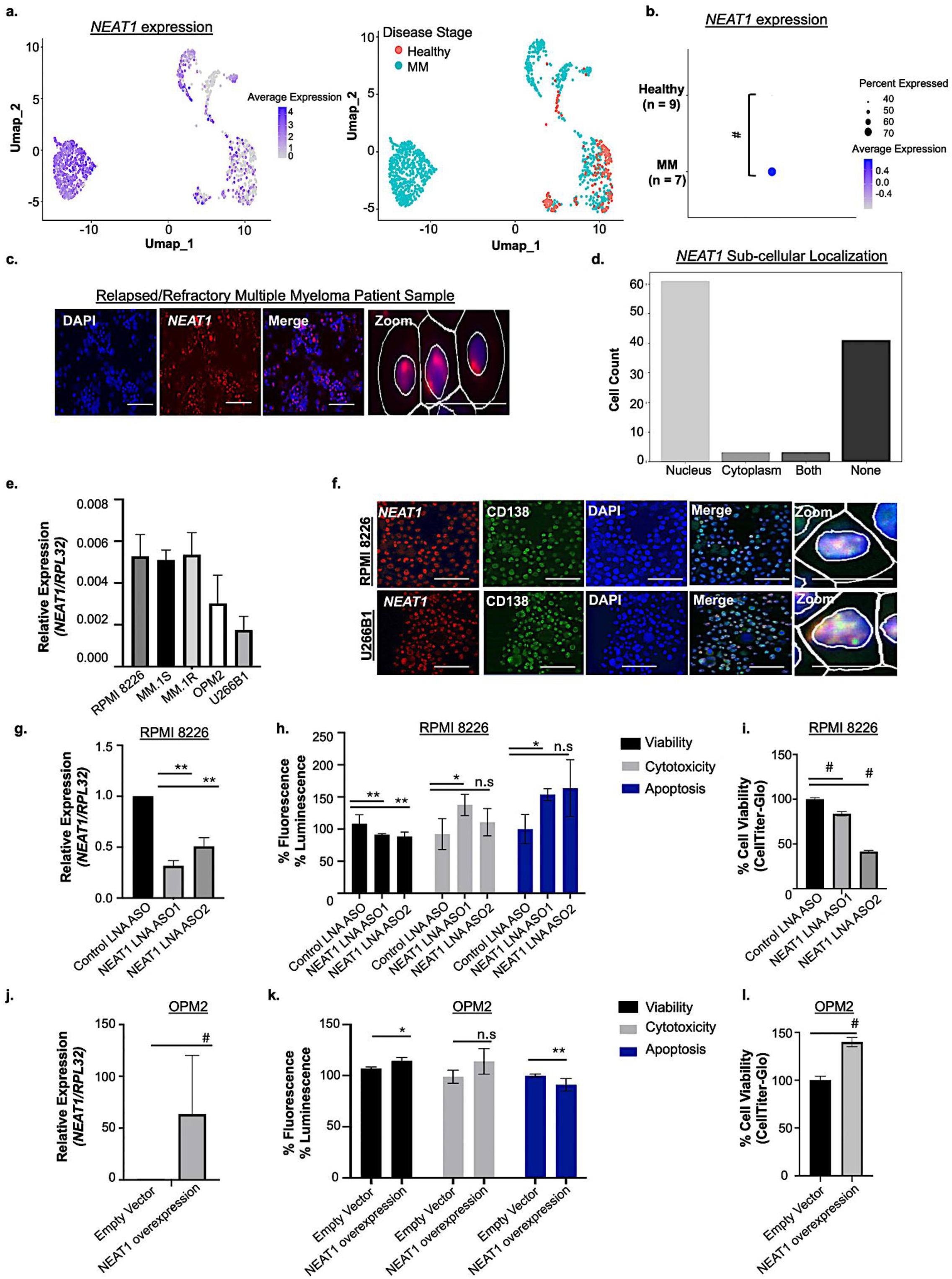
N*E*AT1 is expressed in multiple myeloma and promotes cell viability. **a.** UMAP plot of B and plasma single-cell RNA sequencing data for *NEAT1* (left plot) from healthy (red) and multiple myeloma (MM) patients (blue) (right plot). **b.** Expression of *NEAT1* in MM samples compared to healthy samples. **c.** Multiplex fluorescent in situ hybridization (mFISH) using *NEAT1*-specific probe (red) and DAPI nuclear stain (blue) in relapse/refractory MM (RRMM) patient sample. Scale bar = 20 µM and 60 µM (zoom). **d**. Quantification of subcellular localization of *NEAT1* in RRMM samples from FISH using QuPath analysis. **e.** Expression of *NEAT1* in a panel of MM cell lines using RT-qPCR. **f.** mFISH for *NEAT1* combined with CD138 immunohistochemistry in RPMI 8226 and U266B1 cells. **g.** RT-qPCR quantification of locked nucleic acid (LNA) antisense oligonucleotide (ASO)-mediated knockdown of *NEAT1* in RPMI 8226 cells. **h.** Decreased viability and increased apoptosis in RPMI 8226 cells following LNA ASO-mediated *NEAT1* knockdown, as measured by ApoTox-Glo assay. **i.** Decreased cell viability in RPMI 8226 cells following LNA ASO-mediated *NEAT1* knockdown, as measured by CellTiter-Glo assay. **j.** Expression of *NEAT1* in OPM2 cell lines with *NEAT1* overexpression constructs. **k.** Increased viability and decreased apoptosis in *NEAT1* overexpression OPM2 cells, as measured by ApoTox-Glo assay. **l.** Increased cell viability in *NEAT1* overexpression OPM2 cells, as measured by CellTiter-Glo assay. *p value < 0.05, **p value < 0.005, ***p value < 0.0005, # p value < 0.00005, n.s. not significant.

We next examined *NEAT1* expression in bulk RNA sequencing data from the MMRF CoMMpass study and found that expression was significantly higher in RRMM relative to NDMM disease (p = 0.04; **Supplementary Figure 8b**). Consistent with a lineage-specific role, *NEAT1* expression was also higher in mature B-cell neoplasm cell lines compared to other hematologic malignancies (**Supplementary Figure 8c**). Stratification of RRMM patient samples by genetic subtype further revealed lower *NEAT1* expression in patients with Amp1q, t(11;14), and t(4;14) with higher expression in patients with t(8;14) disease subsets (**Supplementary Figure 8d**).

Next, we optimized detection of *NEAT1* expression in RPMI 8226 cells using multiplexed fluorescent in situ hybridization (mFISH), incorporating a positive control probe (PP1B), a negative control probe (dapB), and *NEAT1*-targeting probes (**Supplementary Figure 9a**). Using this optimized platform, we validated high level expression of *NEAT1* in the nuclear compartment of RRMM patient samples and quantified its subcellular distribution across nuclear, cytoplasmic, dual-localized, and non-expressing cells (54.5% nuclear, 0.045% cytoplasmic, 0.045% both, and 36.3% none, **Figures 6c and d**). We also assessed *NEAT1* expression using RT-qPCR across MM cell lines and observed the highest expression in RPMI 8226, MM.1S, and MM.1R cells, with comparatively lower expression in OPM2 and U266B1 cells (**Figure 6e**). Finally, combined mFISH and immunohistochemistry demonstrated co-localization of *NEAT1* RNA with CD138 protein, a plasma cell-specific marker (**Figure 6f**), and m^6^A (**Supplementary Figure 9b**), confirming *NEAT1* expression within malignant plasma cells.

Together, these analyses establish *NEAT1* as a consistently upregulated lncRNA across MM disease states and genetic subtypes, supporting its functional relevance and motivating investigation of m^6^A-dependent mechanisms of *NEAT1* regulation in MM.

### *NEAT1* knockdown reduces multiple myeloma cell viability and promotes apoptosis

To investigate the functional role of *NEAT1* in MM, we used two independent locked nucleic acid GapmeR antisense oligonucleotides (LNA ASOs) to knock down *NEAT1* in MM cell lines (**Figure 6g**). LNA ASO-mediated *NEAT1* knockdown significantly reduced cell viability in RPMI 8226 cells (**Figures 6h**, LNA ASO1, p = 0.0003; LNA ASO2, p = 0.005; **Figures 6i** LNA ASO1, p = 0.0007; LNA ASO2, p = 1.30 x 10^-06^) and MM.1R cells (LNA ASO1, p = 0.003; LNA ASO2, p = 0.01; **Supplementary Figure 10a**), relative to control LNA ASO. *NEAT1* knockdown also resulted in a modest, but significant increase in apoptosis in RPMI 8226 cells (LNA ASO1, p = 0.004; LNA ASO2, p = 0.06) and MM.1R cells (LNA ASO1, p = 0.03; LNA ASO2, p = 0.006) (**Figures 6h** and Supplementary Figure 10b).

To independently assess *NEAT1* dependency in MM, we interrogated the LongDEP portal, which compiles RNA-targeting CRISPR-Cas13d screens to identify tumor-essential lncRNA isoforms.^49^ In that study, *NEAT1* was identified as a recurrent, isoform-selective cancer dependency across multiple tumor contexts. Consistent with these findings, selective targeting of the *NEAT1-001* isoform using CRISPR-Cas13d resulted in a marked reduction in proliferation across a panel of MM cell lines, including NCI-H929 (log fold change =-3.38), KMS11 (log fold change =-2.87), RPMI 8226 (log fold change =-2.38), OPM2 (log fold change =-1.94), and AMO1 (log fold change =-1.86; **Supplementary Figures 10c and d**). This provides orthogonal validation to the findings of our LNA ASO-based *NEAT1* knockdown experiments. Conversely, *NEAT1* overexpression led to significantly increased viability (p = 0.02 and p = 0.0003; **Figures 6j-l**) and decreased apoptosis (p = 0.003; **Figure 6k**) in OPM2 cells, which express lower endogenous levels of *NEAT1.* Similar increases in viability were observed following *NEAT1* overexpression in U266B1 cells (p = 0.03 and p = 0.04; **Supplementary Figures 11b and c**). Together, these results demonstrate that *NEAT1* promotes MM cell survival and proliferation, consistent with prior reports, and extend these findings by independently validating *NEAT1* dependency using LNA ASOs, supporting a potential therapeutically actionable role for *NEAT1* in MM.

Together, these findings define the RNA modification landscape of MM and identify m^6^A-modified lncRNAs as key components of its regulatory architecture. By integrating transcriptome-wide profiling with site-specific and functional validation, we establish *NEAT1* as a highly m^6^A-modified lncRNA whose expression and activity are regulated by METTL3 (**Figure 7**). Mechanistically, m^6^A modification of *NEAT1*, particularly at site 1611, contributes to MM cell viability and survival, linking epitranscriptomic regulation to oncogenic lncRNA function.

**Figure 7:**
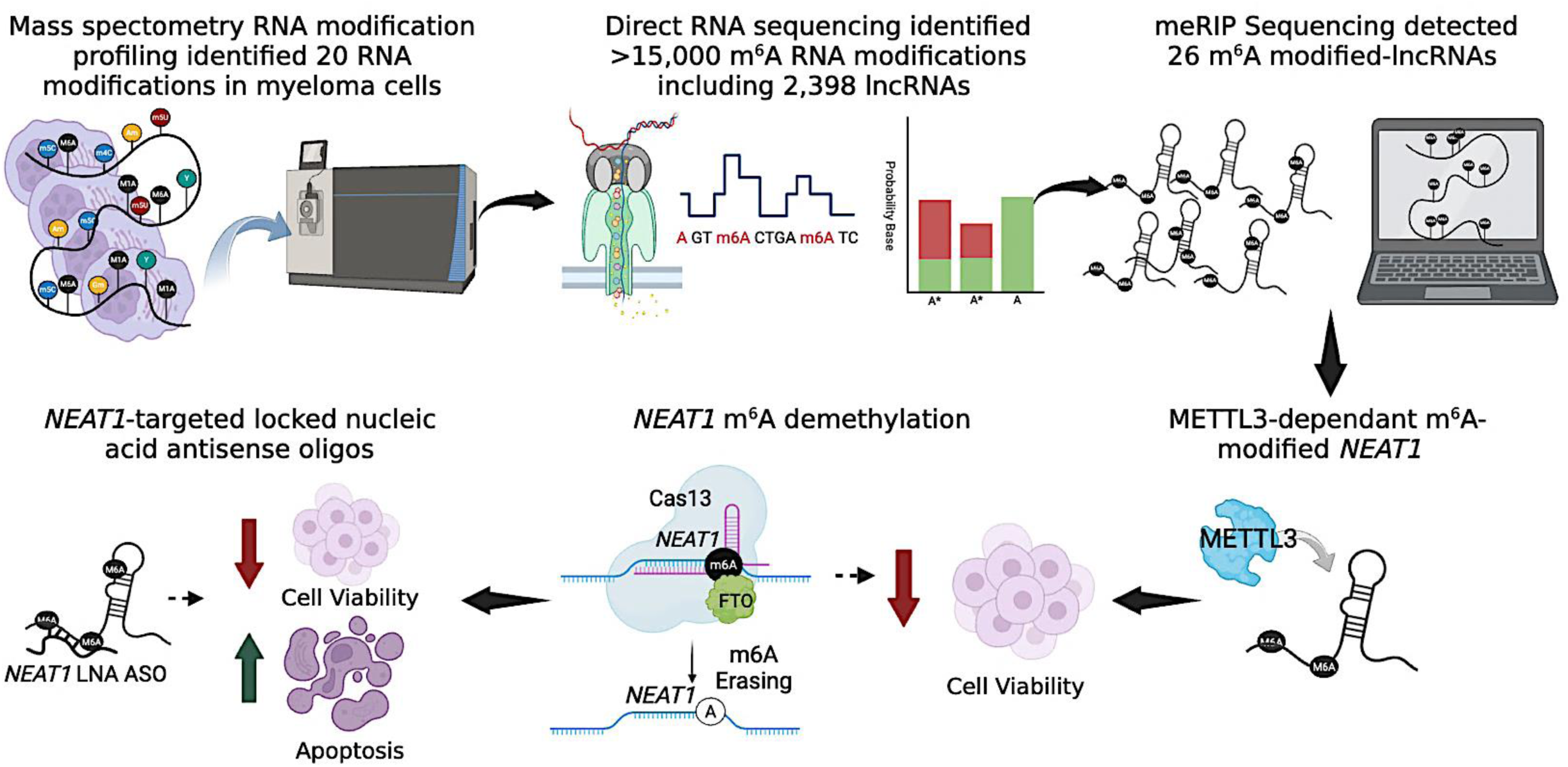
Integrated RNA modification profiling identifies METTL3-dependent m^6^A modification of *NEAT1* as a driver of multiple myeloma. Schematic overview of the experimental strategy and key findings. Global RNA modification profiling of multiple myeloma (MM) cell lines was performed using mass spectrometry-based nucleoside analysis to define the RNA modification landscape in MM, followed by transcriptome-wide m^6^A mapping using direct RNA nanopore sequencing and methylated RNA immunoprecipitation sequencing (meRIP-seq) to identify m^6^A-modified coding and non-coding RNAs. *NEAT1* was identified as a highly m^6^A-modified lncRNA in MM.

## DISCUSSION

MM is an incurable plasma cell malignancy characterized by profound transcriptional plasticity and the inevitable emergence of therapeutic resistance. Despite extensive research into the genetic and epigenetic drivers of MM progression, the contribution of post-transcriptional RNA regulation and RNA modifications remains comparatively underexplored. In this study, we define the RNA modification landscape of MM cell lines, identify m^6^A-modified lncRNAs as key contributors to MM biology, and establish *NEAT1* as a critical epitranscriptomically regulated oncogenic lncRNA whose activity depends on METTL3-mediated m^6^A deposition. By integrating global RNA modification profiling with orthogonal genetic knockdown and pharmacologic inhibition of METTL3 and CRISPR-mediated site-specific RNA demethylation, our findings uncover a previously unappreciated regulatory layer that promotes the survival of MM cells.

m^6^A is the most abundant internal RNA modification in eukaryotic cells, but its functional relevance has been studied predominantly in the context of mRNA metabolism. While lncRNAs are also extensively m^6^A-modified, there is limited functional data linking m^6^A to lncRNA-driven cancer phenotypes, particularly in hematologic malignancies. Our data demonstrate that *NEAT1* is robustly m^6^A-modified in MM and that disruption of this modification, either globally through METTL3 knockdown or inhibition or locally via site-specific demethylation, significantly impairs MM cell viability and promotes apoptosis. A recent review by Dierks et al. proposes that m^6^A functions as a mark of RNA integrity, acting as a selective surveillance mechanism that establishes basal RNA stability rates and potentially additional regulatory functions.^116^ Consistent with this emerging framework, our findings support a model in which m^6^A is not merely a passive modification on lncRNAs, but instead serves as an active regulatory signal required for malignant phenotypes in MM.

Building on this concept, METTL3 has been implicated as an oncogenic factor in MM and other cancers, largely through m^6^A-dependent regulation of mRNA stability, translation, and stemness-associated programs. Our findings extend this paradigm by demonstrating that METTL3 also promotes MM through regulation of oncogenic lncRNAs, specifically *NEAT1*. We show that METTL3 knockdown reduces *NEAT1* expression and MM cell viability, and that pharmacologic inhibition of METTL3 using STM2457 reduces global m^6^A levels and recapitulates these cell viability effects. While additional m^6^A reader proteins such as YTHDF family members or HNRNPA2B1 may also mediate downstream *NEAT1* functions and were not directly interrogated here, the convergence of genetic (siRNA, CRISPR-Cas13d) and chemical inhibition strategies strongly supports on-target disruption of METTL3-dependent RNA methylation as an additional regulatory layer that controls RNA function, in this instance, cell viability in MM cells.

Although previously published bulk RNA-seq analyses showed elevated METTL3 expression comparing healthy and MM plasma cells, this effect was not observed in single-cell RNA-seq analyses of plasma and B-cells comparing healthy and MM plasma cells. This apparent discrepancy suggests that METTL3 upregulation in MM may be governed by enzymatic activity, genetic context (including chromosomal translocations), or microenvironmental cues rather than transcript abundance alone. These observations underscore the importance of functional analyses when assessing epitranscriptomic regulators in cancer.

The demonstration that site-specific removal of m^6^A from *NEAT1* is sufficient to impair MM cell viability without altering overall *NEAT1* expression represents a key conceptual advance of this study. Using a catalytically inactive Cas13a fused to the m^6^A demethylase FTO, we selectively erased m^6^A modifications from a single, highly enriched m^6^A site at position 1611 on *NEAT1*. Despite unchanged *NEAT1* transcript levels, loss of this modification resulted in marked reductions in cell viability and increased apoptosis across MM models. While this study is limited by its focus on a single m^6^A site, rather than combinatorial or multi-site editing, and the absence of direct mechanistic analysis of paraspeckle architecture or transcriptional consequences, these findings still indicate that m^6^A regulates *NEAT1* function rather than transcript stability. This is consistent with emerging models in which m^6^A influences RNA structure, RNA-protein interactions, and subcellular organization. These mechanistic questions are currently under investigation and will be addressed in future studies.

The robustness of our conclusions is supported by orthogonal validation strategies to assess *NEAT1* dependency in MM. In addition to LNA ASO-mediated knockdown, we leveraged publicly available RNA-targeting CRISPR-Cas13d screening data, which independently identified the *NEAT1-001* isoform as essential for proliferation across multiple MM cell lines. Although *in vivo* validation of METTL3 inhibition and *NEAT1* m^6^A editing will be an important next step, the concordance between ASO-mediated depletion, Cas13d-mediated targeting, and site-specific epitranscriptomic editing provides compelling evidence that *NEAT1* is a bona fide dependency in MM. This multi-modal validation mitigates concerns regarding platform-specific artifacts and aligns with prior reports implicating *NEAT1* in MM pathobiology.

Our findings have important therapeutic implications beyond mechanistic insights. METTL3 inhibitors such as STC-15 were originally developed for acute myeloid leukemia, however, our data demonstrate that MM cells are similarly sensitive to METTL3 inhibition, exhibiting reduced m^6^A levels, impaired viability, and increased apoptosis. Several of studies using METTL3 inhibitors are currently being assessed in myeloid malignancies.^46^ Given the emerging role of lncRNAs in therapy resistance and disease persistence, targeting the epitranscriptomic machinery that sustains oncogenic lncRNA function represents an attractive therapeutic strategy. Our data suggest that targeting RNA modifications may disrupt malignant programs without the need to suppress lncRNA expression, potentially offering a therapeutic window with reduced toxicity.

In summary, this study provides an initial framework for understanding the epitranscriptomic landscape of MM and identifies m^6^A-modified *NEAT1* as a critical effector of METTL3-driven oncogenic programs. By demonstrating that site-specific removal of m^6^A is sufficient to impair MM cell survival, we uncover a previously unrecognized mechanism through which RNA modifications regulate lncRNA function in cancer. Together, these findings expand the functional scope of epitranscriptomic regulation in hematologic malignancies and highlight RNA modification machinery as a tractable, mechanistically informed therapeutic vulnerability in MM.

## Acknowledgments

Dr. Jessica Silva-Fisher received funding from the Longer Life Foundation, the Faculty Scholar Award from Washington University Department of Medicine, and Institute of Clinical and Translational Sciences Just-In-time Core usage Funding Program (NIH CTSA Grant #UL1 TR002345). This work is also supported by the Riney Blood Cancer Research Fund. Dr. Benjamin Garcia also received funding from a NIH R01AI118891. Elizabeth Sulvaran-Guel received funding from the American Society of Hematology (Hematology Inclusion Pathway Graduate Award). We would like to thank Dr. John DiPersio for providing cell lines, guidance, and support for this project. We thank the Alvin J. Siteman Cancer Center at Washington University School of Medicine (WUSM) and Barnes-Jewish Hospital in St. Louis, MO, for utilizing the Siteman Flow Cytometry core. We want to thank Dr. Andrew Hutchins from Southern University of Science and Technology for providing the CRISPR-Cas13 plasmids. We also thank Dr. Benjamin Garcia for providing expertise in mass spectrometry and access to machines and data analysis. We appreciate the expertise from Washington University in St. Louis (WashU) Multiple Myeloma Tissue Banking Protocol for sequencing data and access to myeloma tissue samples and the Genome Technology Access Center at the McDonnell Genome Institute at WUSM for processing of Nanopore direct RNA-seq. The Siteman Cancer Center is partly supported by an NCI Cancer Center Support Grant #P30 CA091842. The development of this manuscript was supported by the Scientific Editing Service of the Institute of Clinical and Translational Sciences at WashU, funded by grant UL1TR002345 from the National Center for Advancing Translational Sciences. The content is solely the responsibility of the authors and does not necessarily represent the official view of the NIH.

## Authorship Contributions

P.T. wrote and edited the manuscript, developed cell lines, and conducted *in vitro* assays. D.D. wrote and edited the manuscript, analyzed and processed direct RNA-seq data and mFISH images. C.S. conducted *in vitro* assays. E.S. and S.A. analyzed and processed scRNA-seq data. R.M. conducted *in vitro* assays and mFISH.

J.C conducted in vitro assays. S.G. processed mFISH. S.D., D.C., J.K., C.S. and L.D. conducted and analyzed *in vitro* assays. L.D. and A.K. analyzed *in vitro* data and conducted PubMed searches. K.B. analyzed CLIP-seq data. C.B. conducted mass spectrometry. J.Z., B.G., and J.D. provided project guidance. R.V. and J.D. provided samples. J.S.F. conducted *in vitro* assays, processed mFISH, designed and directed experimental studies, wrote and edited the manuscript, which all authors reviewed and approved.

---

**Supplementary Figure 1:**
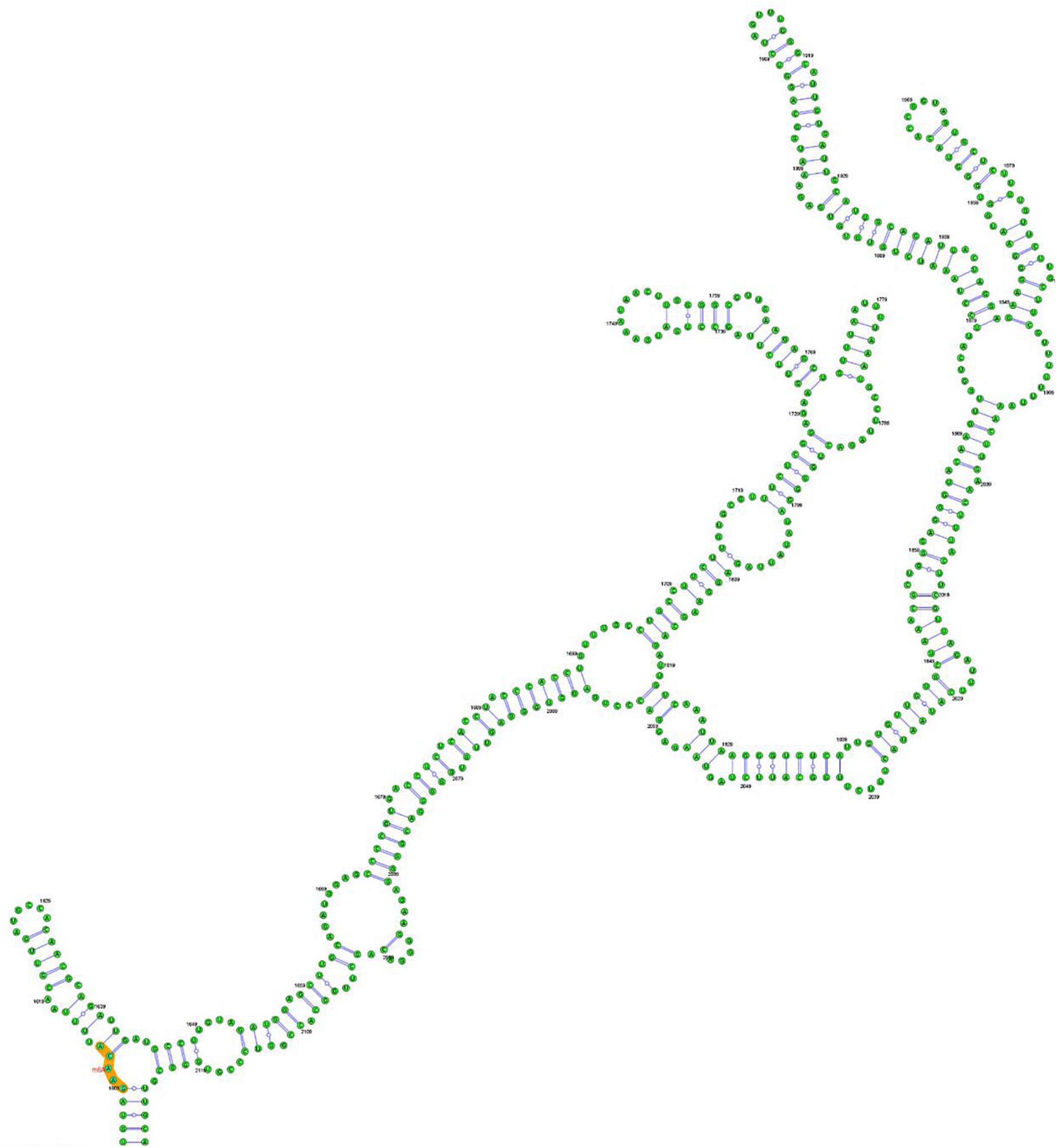
Structure of *NEAT1* m^6^A 1611 site. RNA structure of *NEAT1* highlighting m6A 1611 site (orange) using sequence-basedRNAadenosinemethylation sitepredictor site.

**Supplementary Figure 2:**
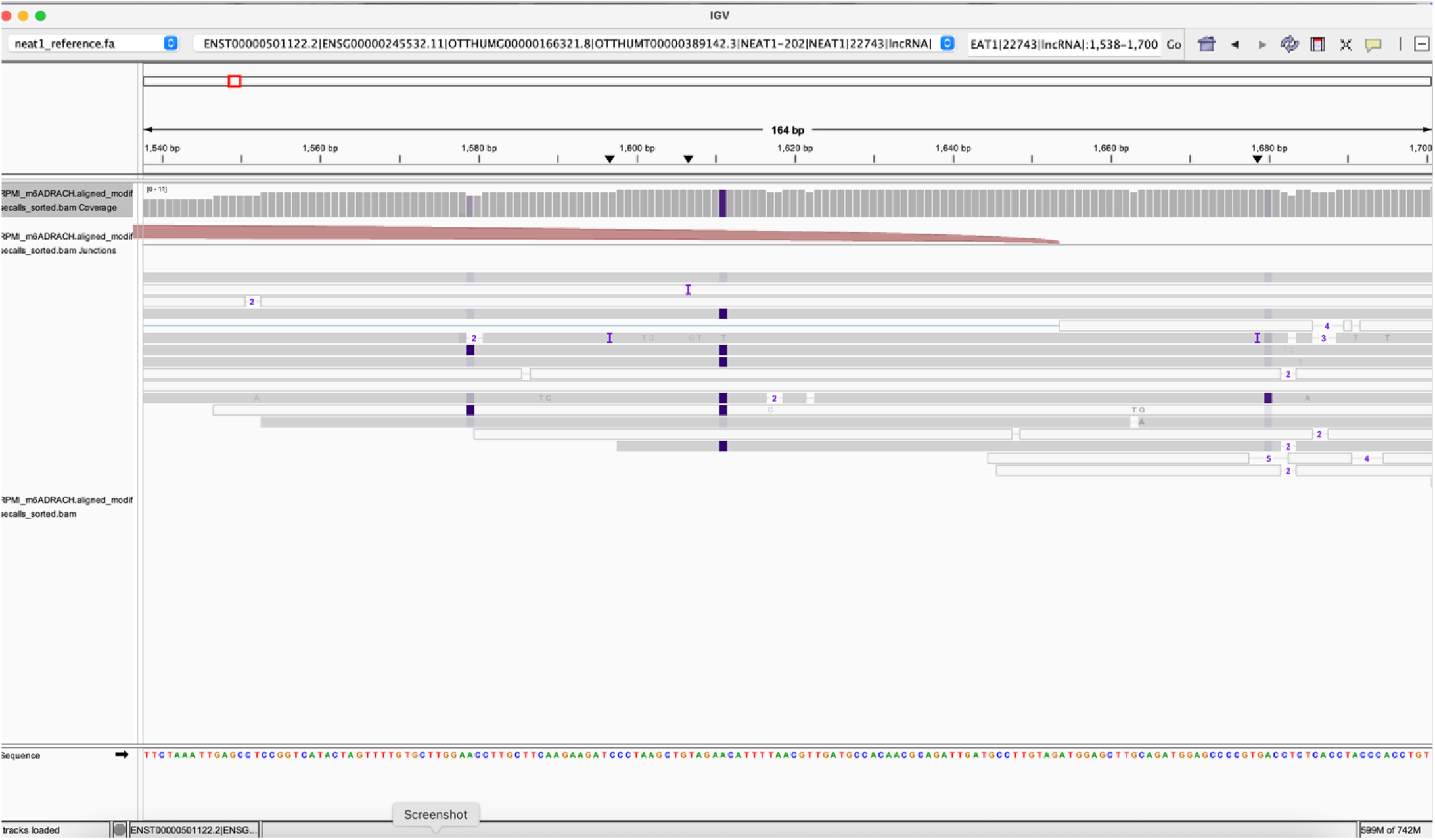
m^6^A identified peaks and *NEAT1* m6A peak at site 1611 in RPMI 8226 cells. IGV viewer showing *NEAT1* m^6^A 1611 peak in RPMI 8226 cells detected using Nanopore Direct-Sequencing. Purple indicates m^6^A coverage.

**Supplementary Figure 3:**
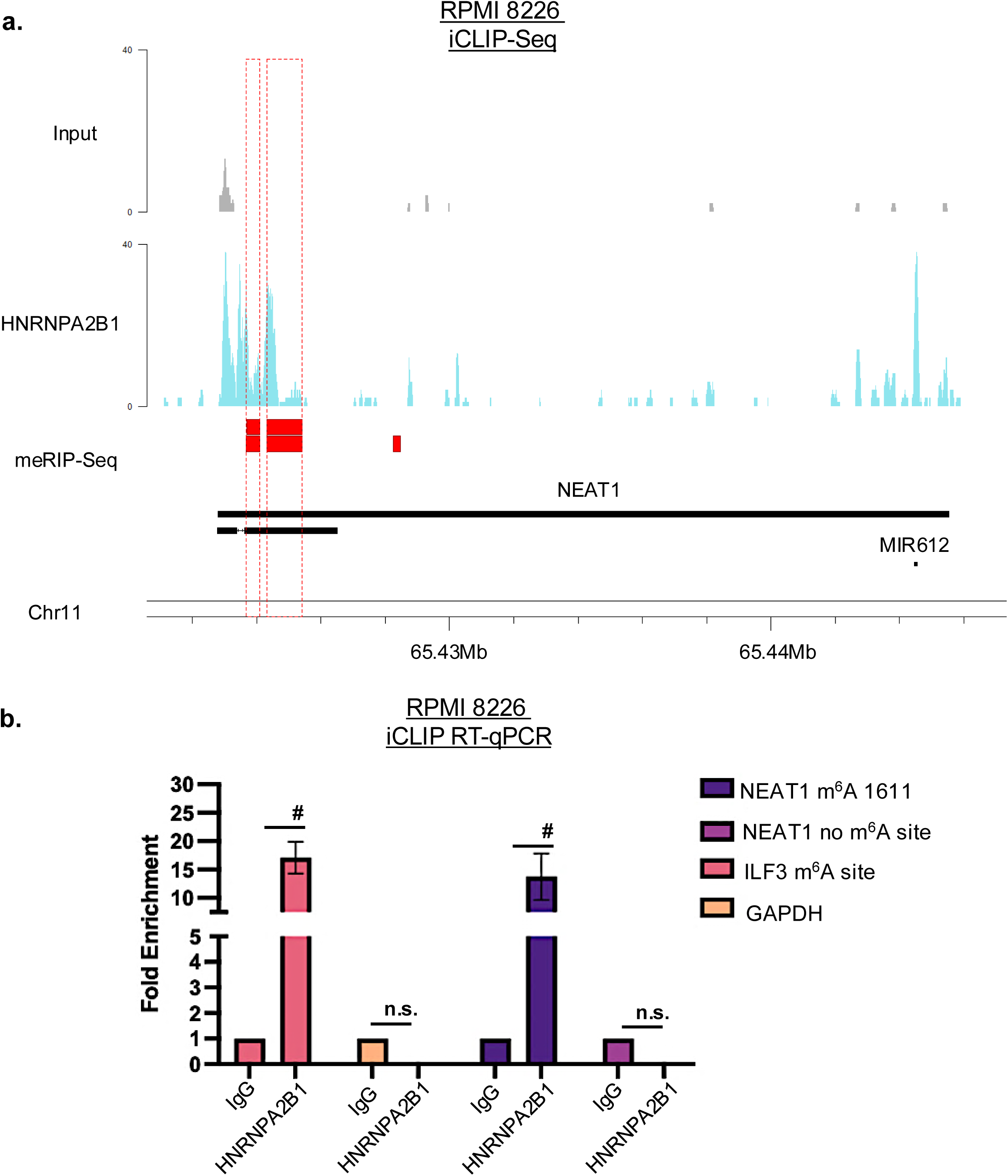
*NEAT1* enrichment peaks for HNRNPA2B1 CLIP-Seq. a. IGV viewer showing *NEAT1* enrichment peaks in HNRNA2B1 UV crosslinking immunoprecipitation sequencing (CLIP-Seq) compared to Input and b. HNRNPA2B1 CLIP RT-qPCR showing *NEAT1* high fold enrichment for *NEAT1* m^6^A site 1611. #p value < 0.0005

**Supplementary Figure 4:**
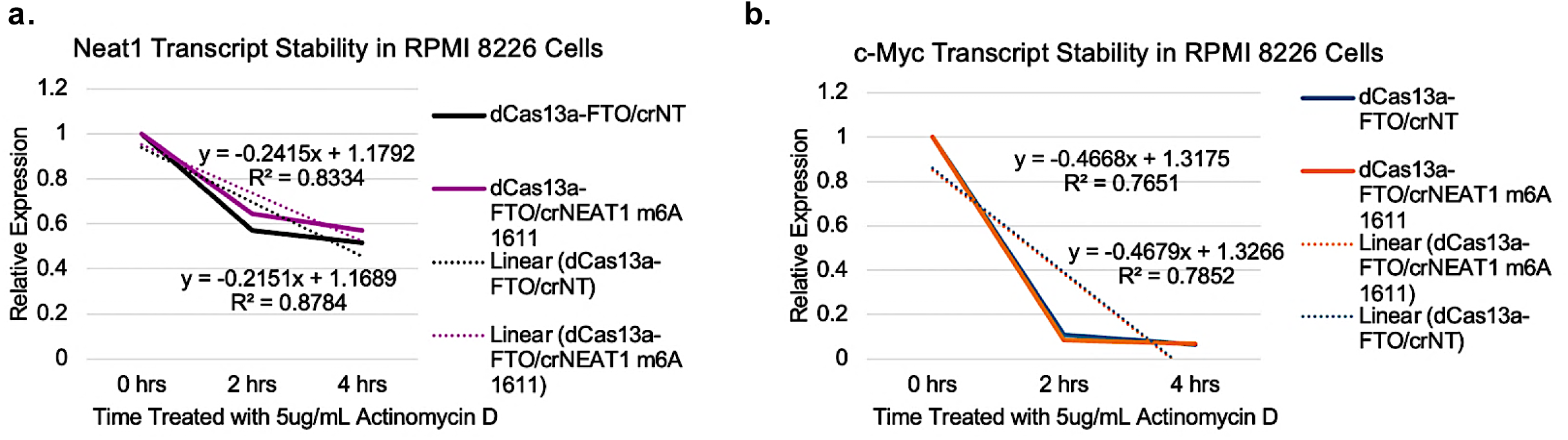
*NEAT1* m^6^A 1611 site and stability of *NEAT1* transcript. RPMI 8226 cell line treated with 5ug/ul Actinomycin D for increasing concentrations showing expression of a. *NEAT1* and b. *c-Myc*.

**Supplementary Figure 5:**
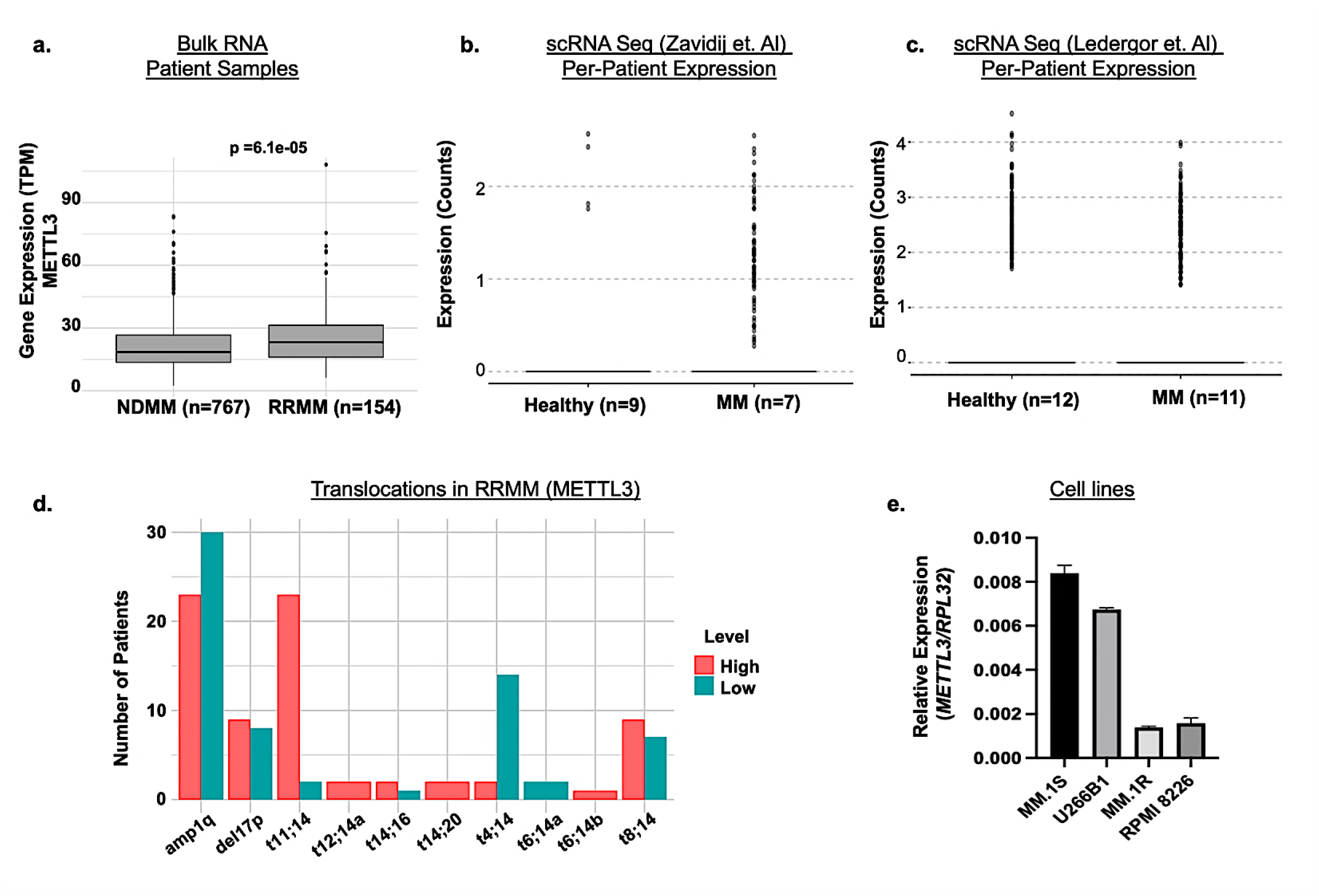
RNA expression and genetic translocations of *METTL3* in myeloma cell lines. **a.** RNA sequencing expression of *METTL3* in newly diagnosed multiple myeloma (MM) patient samples and relapsed/refractory MM samples. **b.** and **c.** Expression of *METTL3* in B cells and Plasma cells from healthy and MM samples using single-cell RNA sequencing dot plot. **d.** Molecular characterization of *METTL3* in relapsed/refractory MM patient data. **e.** RT-qPCR expression of *METTL3* in multiple myeloma cell line panel.

**Supplementary Figure 6:**
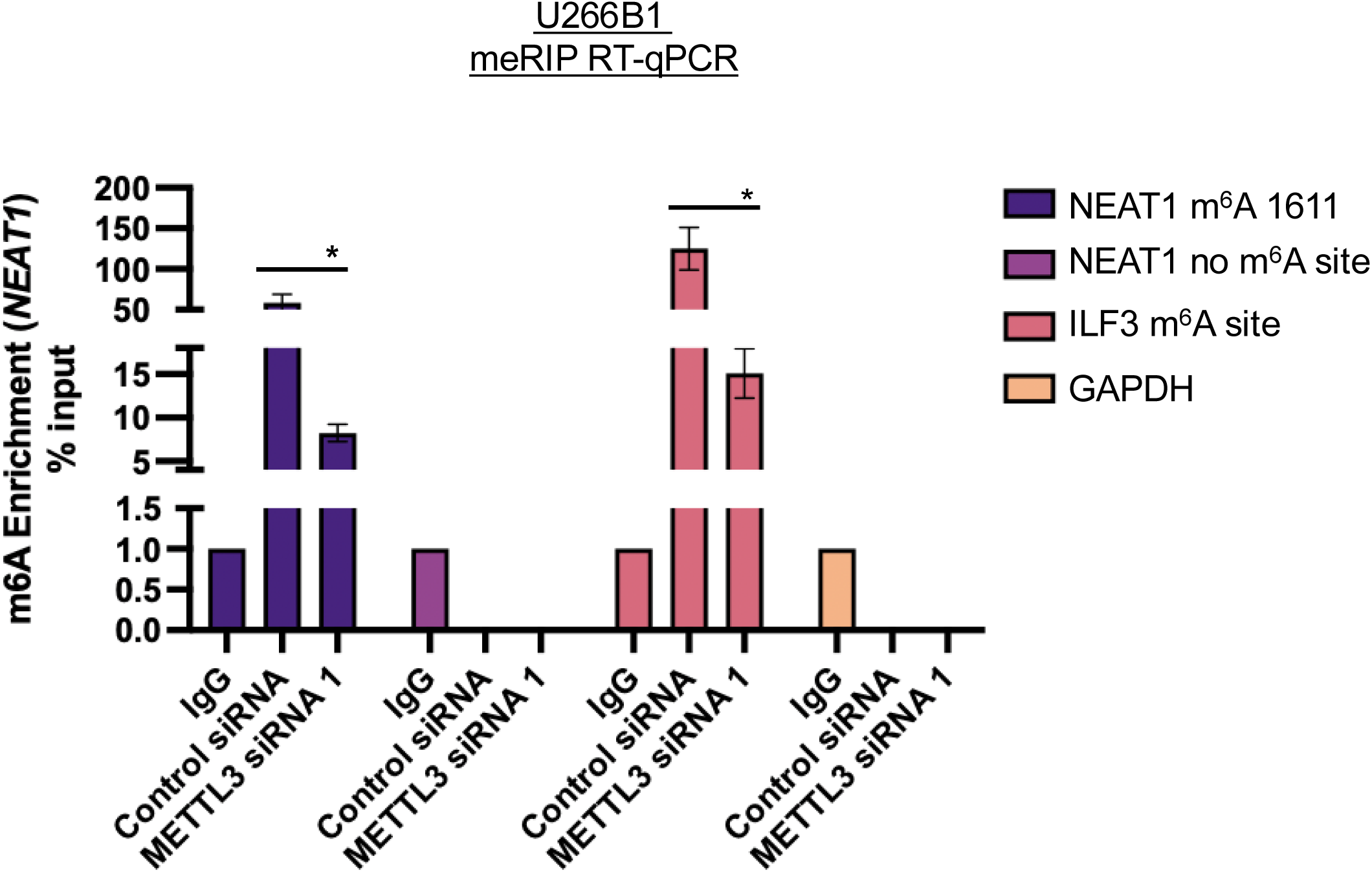
***METTL3* knockdown decreases m6A modification on *NEAT1*** U266B1 meRIP RT-qPCR showing *NEAT1* high m6A fold enrichment for *NEAT1* m^6^A site 1611 with significant decrease of m^6^A when compared to knockdown with METTL3 silencer RNA (siRNA). *p value < 0.05

**Supplementary Figure 7:**
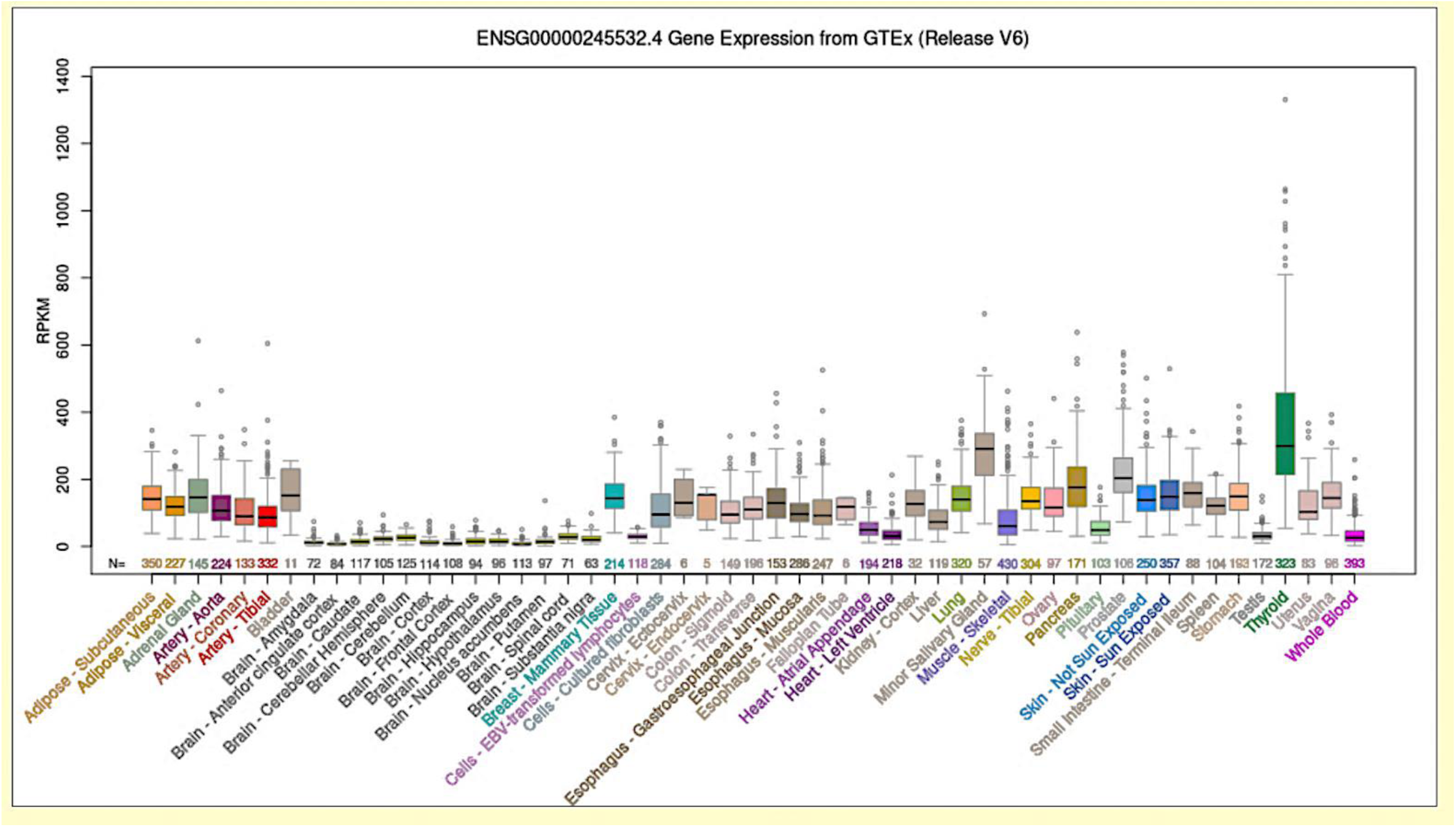
*NEAT1* expression in normal tissues. *NEAT1* gene expression in a panel of 53 normal tissues from the Genotype-Tissue Expression (GTEx) Portal Release V6, data from UCSC Browser.

**Supplementary Figure 8:**
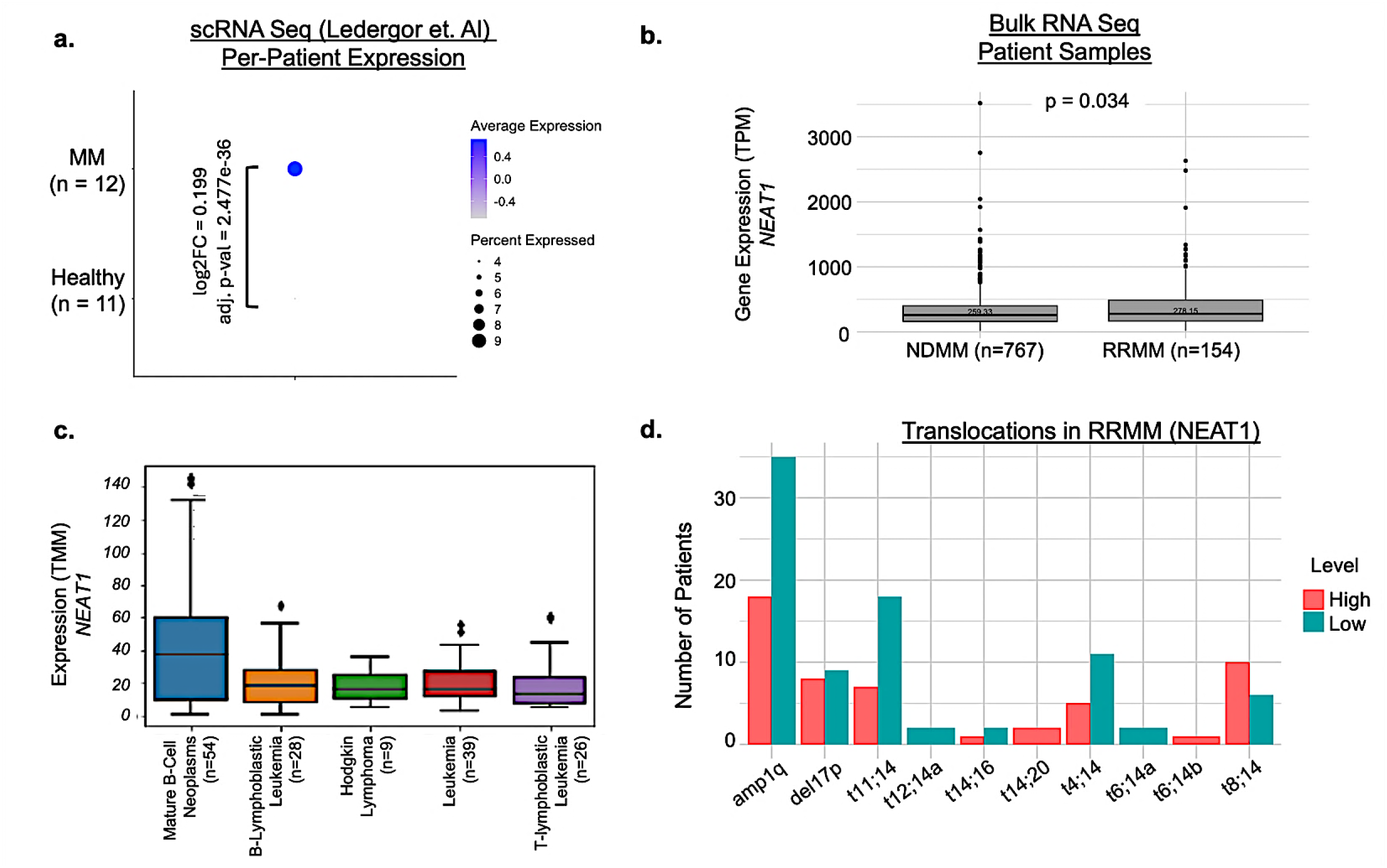
Expression of *NEAT1* in blood cancer and translocations. **a.** Expression of *NEAT1* in B cells and Plasma cells from healthy and multiple myeloma (MM) samples using single-cell RNA sequencing dot plot. **b.** RNA sequencing expression of *NEAT1* in newly diagnosed MM patient samples and relapsed/refractory MM samples and **c.** blood cancer cell lines from cBioPortal. **d.** Molecular characterization of *NEAT1* in relapsed/refractory MM patient data.

**Supplementary Figure 9:**
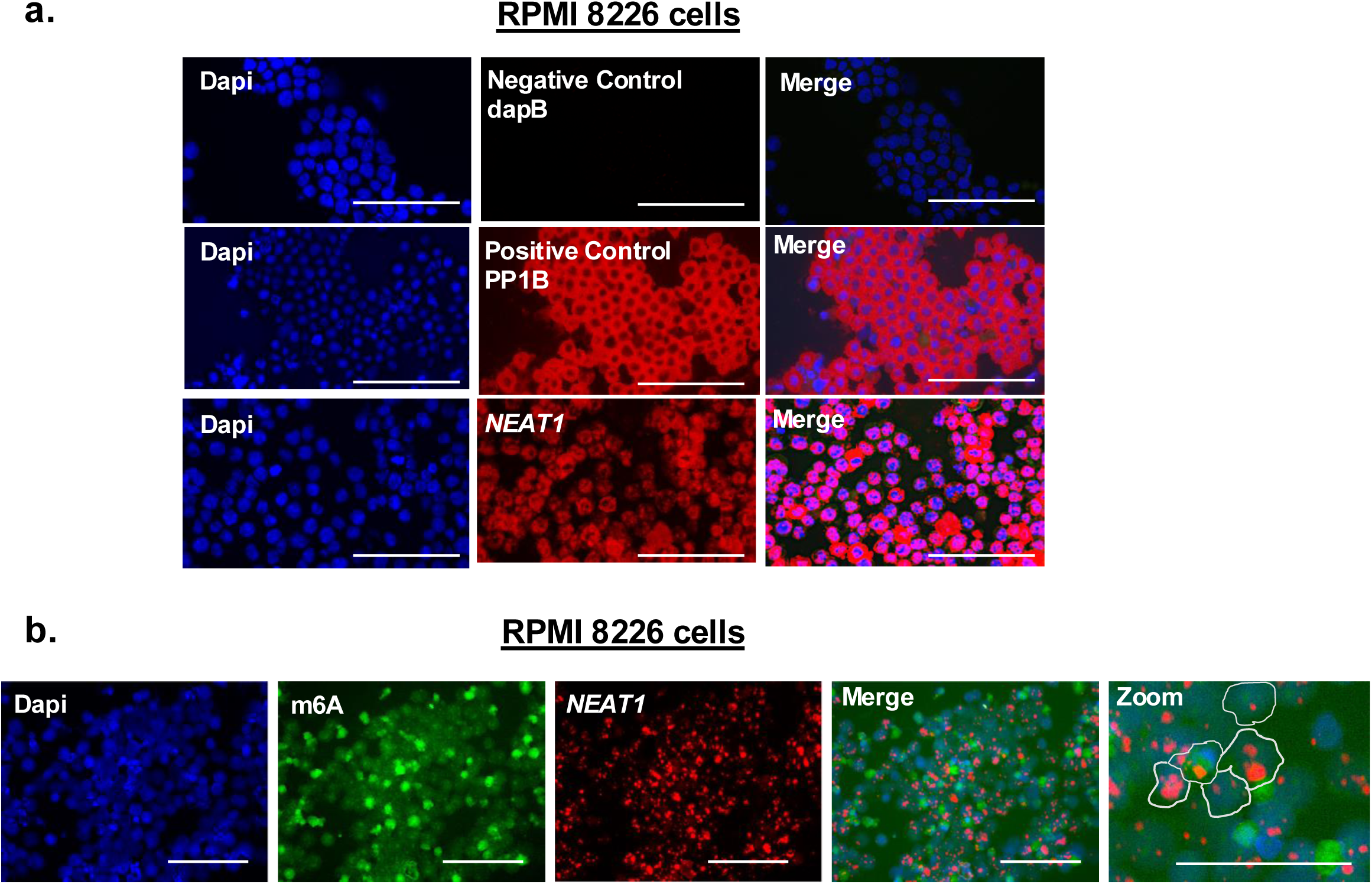
Detection of *NEAT1* and m6A using multiplexed RNA in situ hybridization in myeloma cells. **a.** Expression of negative control, positive control, and *NEAT1* targeted probes for multiplexed RNA in situ hybridization in RPMI 8226 myeloma cells and **b.** co-hybridization using m^6^A antibody. Dapi (nucleus; blue), Targets (dapB, PPIB, *NEAT1*; red), Scale bars 20x and 40x

**Supplementary Figure 10:**
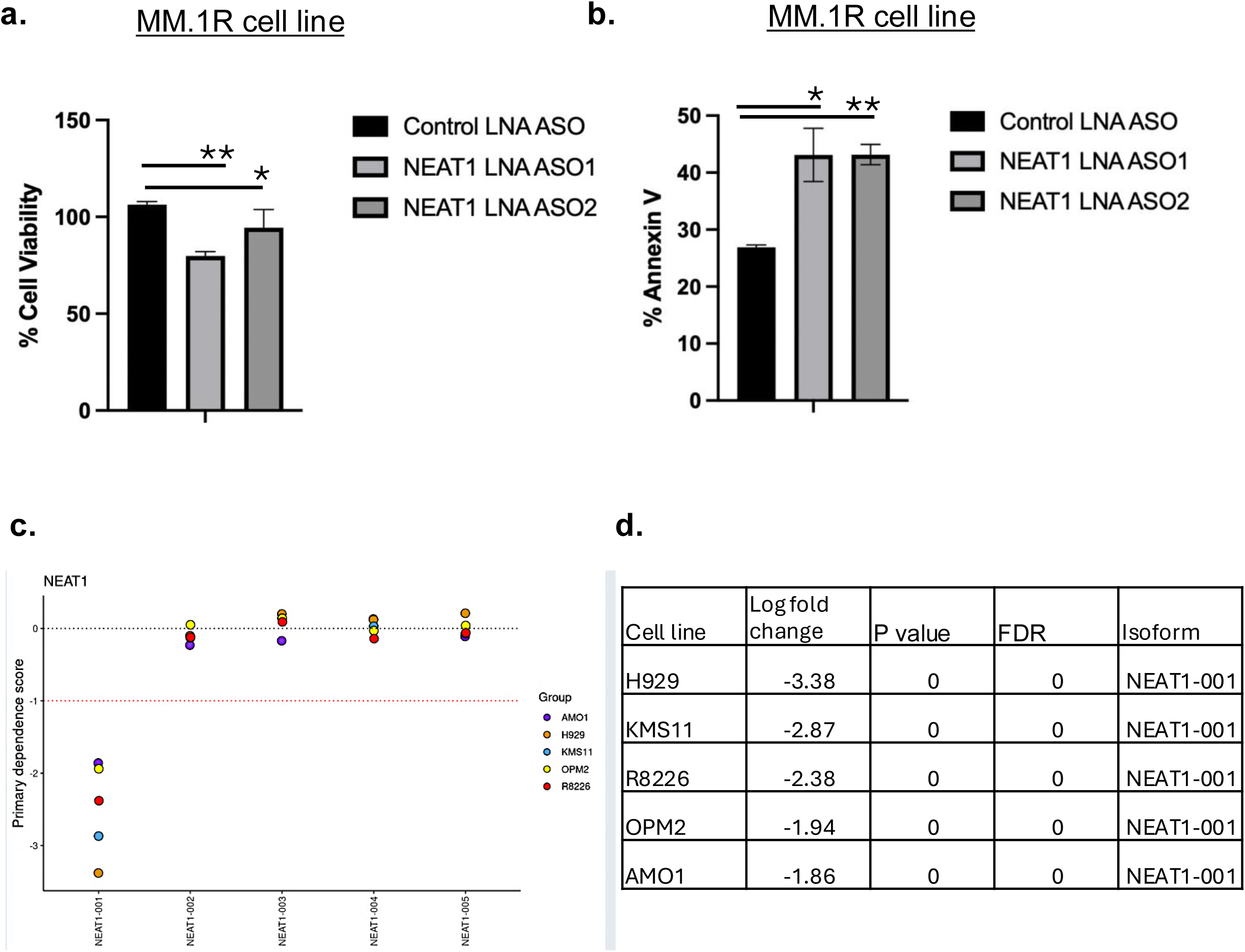
Knockdown of *NEAT1* decreases proliferation and increases apoptosis. **a.** RT-qPCR quantification of locked nucleic acid (LNA) antisense oligonucleotide (ASO)-mediated knockdown of *NEAT1* in MM.1R cells. **b.** MM.1R cells showing increased apoptosis following *NEAT1* knockdown using Annexin V assay. **c.** CRISPR-Cas13a screen for *NEAT1-001* isoform showing decreased viability in panel of myeloma cell lines and **d**. associated statistics from LongDEP Portal. *p value < 0.05, **p value < 0.005

**Supplementary Figure 11:**
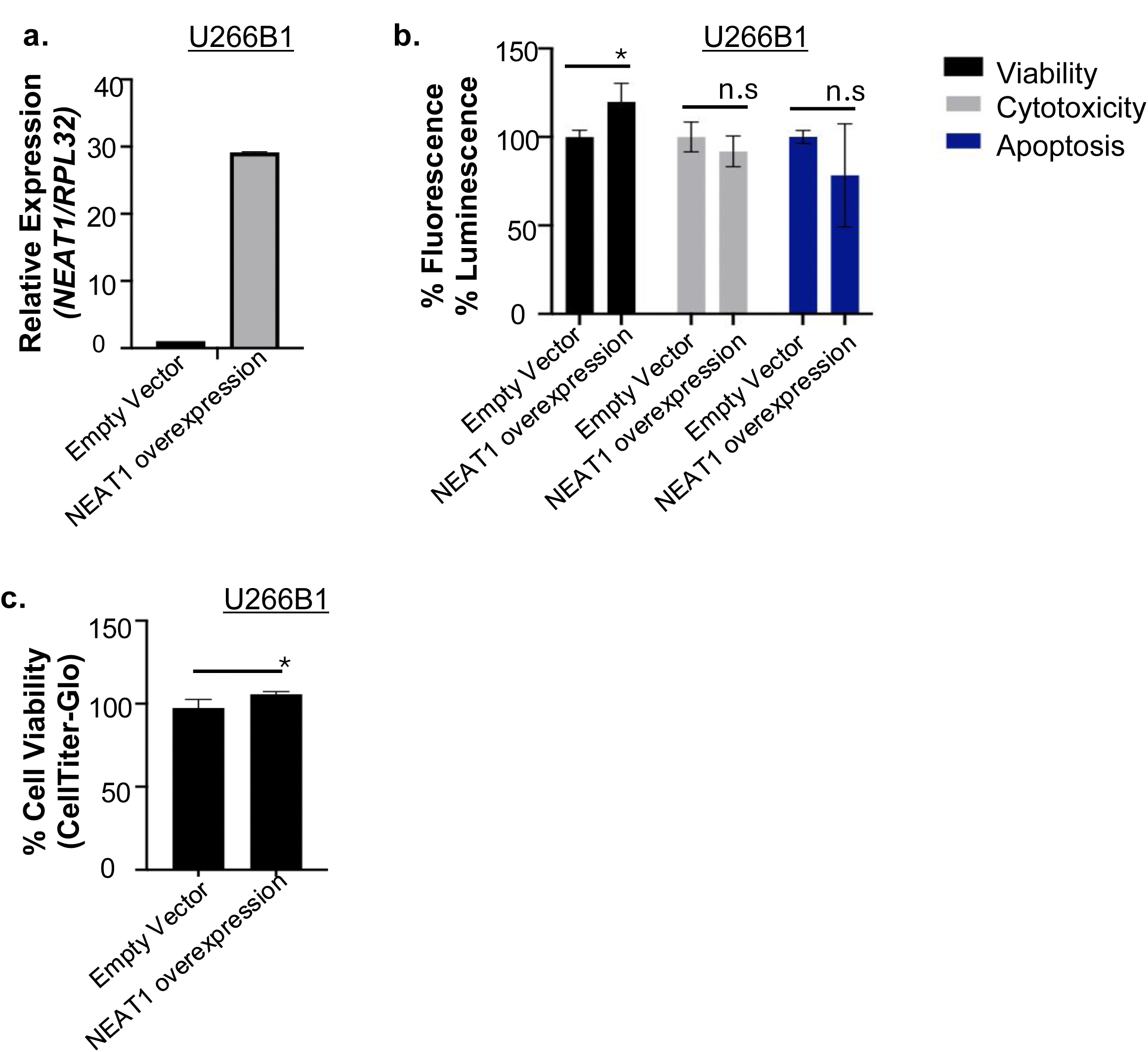
Overexpressing *NEAT1* in U266B1 cells increases proliferation. a. RT-qPCR quantification of *NEAT1* in U266B1 cells overexpressing *NEAT1*. b. U266B1 overexpression cells showing increased viability following *NEAT1* overexpression using ApoTox-Glo assay and c. CellTiter-Glo assay. *p value < 0.05

## BIBLIOGRAPHY

1. Wu, Y., Zheng, Y.Y., Lin, Q. & Sheng, J. Detection and Quantification of RNA Phosphorothioate Modifications Using Mass Spectrometry. Curr Protoc Nucleic Acid Chem 82, e113 (2020).

2. Janssen, K.A., Xie, Y., Kramer, M.C., Gregory, B.D. & Garcia, B.A. Data-Independent Acquisition for the Detection of Mononucleoside RNA Modifications by Mass Spectrometry. J Am Soc Mass Spectrom 33, 885–893 (2022).

3. Ron, K., Kahn, J., Malka-Tunitsky, N. & Sas-Chen, A. High-throughput detection of RNA modifications at single base resolution. FEBS Lett 599, 19–32 (2025).

4. Yuan, X., et al. Mass Spectrometry-Based Direct Sequencing of tRNAs De Novo and Quantitative Mapping of Multiple RNA Modifications. J Am Chem Soc 146, 25600–25613 (2024).

5. Hermon, S.J., Sennikova, A. & Becker, S. Quantitative detection of pseudouridine in RNA by mass spectrometry. Scientific reports 14, 27564 (2024).

6. Smith, M.A., et al. Molecular barcoding of native RNAs using nanopore sequencing and deep learning. Genome Res 30, 1345–1353 (2020).

7. Liu, H., et al. Accurate detection of m(6)A RNA modifications in native RNA sequences. Nature communications 10, 4079 (2019).

8. Mulroney, L., Birney, E., Leonardi, T. & Nicassio, F. Using Nanocompore to Identify RNA Modifications from Direct RNA Nanopore Sequencing Data. Curr Protoc 3, e683 (2023).

9. Wu, Z., et al. Nanopore direct RNA sequencing for RNA modification analysis: workflow assessment and computational tool benchmarking. Adv Biotechnol (Singap) 4(2026).

10. Leger, A., et al. RNA modifications detection by comparative Nanopore direct RNA sequencing. Nature communications 12, 7198 (2021).

11. Dominissini, D., et al. Topology of the human and mouse m6A RNA methylomes revealed by m6A-seq. Nature 485, 201–206 (2012).

12. Meyer, K.D., et al. Comprehensive analysis of mRNA methylation reveals enrichment in 3’ UTRs and near stop codons. Cell 149, 1635–1646 (2012).

13. Saletore, Y., et al. The birth of the Epitranscriptome: deciphering the function of RNA modifications. Genome Biol 13, 175 (2012).

14. Jonkhout, N., et al. The RNA modification landscape in human disease. Rna 23, 1754–1769 (2017).

15. Wein, S., et al. A computational platform for high-throughput analysis of RNA sequences and modifications by mass spectrometry. Nature communications 11, 926 (2020).

16. Xie, Y., et al. Permethylation of Ribonucleosides Provides Enhanced Mass Spectrometry Quantification of Post-Transcriptional RNA Modifications. Anal Chem 94, 7246–7254 (2022).

17. Warneford-Thomson, R., He, C., Sidoli, S., Garcia, B.A. & Bonasio, R. Sample Preparation for Mass Spectrometry-based Identification of RNA-binding Regions. J Vis Exp (2017).

18. Krakau, S., Richard, H. & Marsico, A. PureCLIP: capturing target-specific protein-RNA interaction footprints from single-nucleotide CLIP-seq data. Genome Biol 18, 240 (2017).

19. Van Nostrand, E.L., et al. Robust transcriptome-wide discovery of RNA-binding protein binding sites with enhanced CLIP (eCLIP). Nat Methods 13, 508–514 (2016).

20. Guo, T., Liu, D.F., Peng, S.H. & Xu, A.M. ALKBH5 promotes colon cancer progression by decreasing methylation of the lncRNA NEAT1. Am J Transl Res 12, 4542–4549 (2020).

21. Liu, J., Zhao, W., Zhang, L. & Wang, X. The emerging roles of N6-methyladenosine (m6A)-modified long non-coding RNAs in human cancers. Cell death discovery 8, 255 (2022).

22. Yi, Y.C., Chen, X.Y., Zhang, J. & Zhu, J.S. Novel insights into the interplay between m(6)A modification and noncoding RNAs in cancer. Mol Cancer 19, 121 (2020).

23. Zhang, S., et al. m(6)A Demethylase ALKBH5 Maintains Tumorigenicity of Glioblastoma Stem-like Cells by Sustaining FOXM1 Expression and Cell Proliferation Program. Cancer cell 31, 591–606 e596 (2017).

24. Huang, Y., et al. Small-Molecule Targeting of Oncogenic FTO Demethylase in Acute Myeloid Leukemia. Cancer cell 35, 677–691 e610 (2019).

25. Lin, S., Choe, J., Du, P., Triboulet, R. & Gregory, R.I. The m(6)A Methyltransferase METTL3 Promotes Translation in Human Cancer Cells. Molecular cell 62, 335–345 (2016).

26. Vu, L.P., et al. The N(6)-methyladenosine (m(6)A)-forming enzyme METTL3 controls myeloid differentiation of normal hematopoietic and leukemia cells. Nature medicine 23, 1369–1376 (2017).

27. Barbieri, I. & Kouzarides, T. Role of RNA modifications in cancer. Nat Rev Cancer 20, 303–322 (2020).

28. Qin, X., et al. RNA modifications in cancer immune therapy: regulators of immune cells and immune checkpoints. Front Immunol 15, 1463847 (2024).

29. Huang, G., Qiu, Y., Fan, Y. & Liu, J. METTL3-deficiency Suppresses Neural Apoptosis to Induce Protective Effects in Cerebral I/R Injury via Inhibiting RNA m6A Modifications: A Pre-clinical and Pilot Study. Neurochem Res 49, 85–98 (2024).

30. Dutheuil, G., et al. Discovery, Optimization, and Preclinical Pharmacology of EP652, a METTL3 Inhibitor with Efficacy in Liquid and Solid Tumor Models. J Med Chem 68, 2981–3003 (2025).

31. Guirguis, A.A., et al. Inhibition of METTL3 Results in a Cell-Intrinsic Interferon Response That Enhances Antitumor Immunity. Cancer Discov 13, 2228–2247 (2023).

32. Huarte, M. The emerging role of lncRNAs in cancer. Nature medicine 21, 1253–1261 (2015).

33. Silva, J. & Smith, D. Long non-coding RNAs and Cancer, (Caister Academic Press, La Jolla, California, 2012).

34. Mattick, J.S., et al. Long non-coding RNAs: definitions, functions, challenges and recommendations. Nature reviews. Molecular cell biology 24, 430–447 (2023).

35. Kopp, F. & Mendell, J.T. Functional Classification and Experimental Dissection of Long Noncoding RNAs. Cell 172, 393–407 (2018).

36. Herman, A.B., Tsitsipatis, D. & Gorospe, M. Integrated lncRNA function upon genomic and epigenomic regulation. Molecular cell 82, 2252–2266 (2022).

37. Statello, L., Guo, C.J., Chen, L.L. & Huarte, M. Author Correction: Gene regulation by long non-coding RNAs and its biological functions. Nature reviews. Molecular cell biology 22, 159 (2021).

38. He, R.Z., Jiang, J. & Luo, D.X. The functions of N6-methyladenosine modification in lncRNAs. Genes Dis 7, 598–605 (2020).

39. Wang, Z.W., et al. SRSF3-mediated regulation of N6-methyladenosine modification-related lncRNA ANRIL splicing promotes resistance of pancreatic cancer to gemcitabine. Cell reports 39, 110813 (2022).

40. Siegel, R.L., Giaquinto, A.N. & Jemal, A. Cancer statistics, 2024. CA: a cancer journal for clinicians 74, 12–49 (2024).

41. Gajek, A., et al. Chemical modification of melphalan as a key to improving treatment of haematological malignancies. Scientific reports 10, 4479 (2020).

42. Lee, J.H. & Kim, S.H. Treatment of relapsed and refractory multiple myeloma. Blood Res 55, S43–S53 (2020).

43. Wang, G., et al. N6-methyladenosine-mediated upregulation of H19 promotes resistance to bortezomib by modulating the miR-184/CARM1 axis in multiple myeloma. Clin Exp Med 25, 102 (2025).

44. Bao, J., et al. N6-methyladenosine-induced miR-182-5p promotes multiple myeloma tumorigenesis by regulating CAMK2N1. Mol Cell Biochem 479, 3077–3089 (2024).

45. Lu, X., et al. Regulatory role of the METTL3/MALAT1 axis in multiple myeloma progression. J Bone Oncol 53, 100695 (2025).

46. Huang, X., Yang, Z., Li, Y. & Long, X. m6A methyltransferase METTL3 facilitates multiple myeloma cell growth through the m6A modification of BZW2. Ann Hematol 102, 1801–1810 (2023).

47. Che, F., et al. METTL3 facilitates multiple myeloma tumorigenesis by enhancing YY1 stability and pri-microRNA-27 maturation in m(6)A-dependent manner. Cell Biol Toxicol 39, 2033–2050 (2023).

48. Chen, C.J., et al. Metformin attenuates multiple myeloma cell proliferation and encourages apoptosis by suppressing METTL3-mediated m6A methylation of THRAP3, RBM25, and USP4. Cell Cycle 22, 986–1004 (2023).

49. Morelli, E., et al. CRISPR-Cas13d functional transcriptomics reveals widespread isoform-selective cancer dependencies on lncRNAs. Blood 146, 847–860 (2025).

50. Mishra, R., et al. Investigation of lncRNA expression in newly diagnosed multiple myeloma reveals a LINC01432-CELF2 axis as an inhibitor of apoptosis. Oncogenesis 14, 36 (2025).

51. Ronchetti, D., et al. A compendium of long non-coding RNAs transcriptional fingerprint in multiple myeloma. Scientific reports 8, 6557 (2018).

52. Carrasco-Leon, A., et al. Characterization of complete lncRNAs transcriptome reveals the functional and clinical impact of lncRNAs in multiple myeloma. Leukemia 35, 1438–1450 (2021).

53. Zhou, M., et al. Identification and validation of potential prognostic lncRNA biomarkers for predicting survival in patients with multiple myeloma. J Exp Clin Cancer Res 34, 102 (2015).

54. Zavidij, O., et al. Single-cell RNA sequencing reveals compromised immune microenvironment in precursor stages of multiple myeloma. Nat Cancer 1, 493–506 (2020).

55. Ledergor, G., et al. Single cell dissection of plasma cell heterogeneity in symptomatic and asymptomatic myeloma. Nature medicine 24, 1867–1876 (2018).

56. Jiang, F., et al. HNRNPA2B1 promotes multiple myeloma progression by increasing AKT3 expression via m6A-dependent stabilization of ILF3 mRNA. Journal of hematology & oncology 14, 54 (2021).

57. Puram, S.V., et al. Cellular states are coupled to genomic and viral heterogeneity in HPV-related oropharyngeal carcinoma. Nat Genet 55, 640–650 (2023).

58. Chang, C., Ma, G., Cheung, E. & Hutchins, A.P. A programmable system to methylate and demethylate N(6)-methyladenosine (m(6)A) on specific RNA transcripts in mammalian cells. J Biol Chem 298, 102525 (2022).

59. Yuan, Y., Du, Y., Wang, L. & Liu, X. The M6A methyltransferase METTL3 promotes the development and progression of prostate carcinoma via mediating MYC methylation. J Cancer 11, 3588–3595 (2020).

60. Ma, Y., Shi, H. & Zheng, W. METTL3 Regulates the Translation of Oncogene Myc through m(6)A Modification and Promotes the Occurrence and Development of Cervical Cancer. Discov Med 36, 1902–1910 (2024).

61. Wang, F., et al. LncRNA FTO-IT1 promotes glycolysis and progression of hepatocellular carcinoma through modulating FTO-mediated N6-methyladenosine modification on GLUT1 and PKM2. J Exp Clin Cancer Res 42, 267 (2023).

62. Edupuganti, R.R., et al. N(6)-methyladenosine (m(6)A) recruits and repels proteins to regulate mRNA homeostasis. Nature structural & molecular biology 24, 870–878 (2017).

63. Han, Y., Sun, J., Yao, M., Miao, L. & Li, M. Biological roles of enhancer RNA m6A modification and its implications in cancer. Cell Commun Signal 23, 254 (2025).

64. Mao, Y., et al. METTL3-Mediated m(6)A Modification of lncRNA MALAT1 Facilitates Prostate Cancer Growth by Activation of PI3K/AKT Signaling. Cell Transplant 31, 9636897221122997 (2022).

65. Tian, Y., Xiao, Y.H., Sun, C., Liu, B. & Sun, F. N6-Methyladenosine Methyltransferase METTL3 Alleviates Diabetes-Induced Testicular Damage through Modulating TUG1/Clusterin Axis. Diabetes Metab J 47, 287–300 (2023).

66. Yao, F.Y., et al. m(6)A Modification of lncRNA NEAT1 Regulates Chronic Myelocytic Leukemia Progression via miR-766-5p/CDKN1A Axis. Frontiers in oncology 11, 679634 (2021).

67. Wen, S., et al. Long non-coding RNA NEAT1 promotes bone metastasis of prostate cancer through N6-methyladenosine. Mol Cancer 19, 171 (2020).

68. Alarcon, C.R., et al. HNRNPA2B1 Is a Mediator of m(6)A-Dependent Nuclear RNA Processing Events. Cell 162, 1299–1308 (2015).

69. Jia, C., et al. HNRNPA2B1-mediated m6A modification of TLR4 mRNA promotes progression of multiple myeloma. Journal of translational medicine 20, 537 (2022).

70. Chen, J., et al. Targeted Methylation of the LncRNA NEAT1 Suppresses Malignancy of Renal Cell Carcinoma. Front Cell Dev Biol 9, 777349 (2021).

71. Chen, Q., et al. Prognostic genes related to mitochondrial dynamics and mitophagy in diffuse large B-cell lymphoma are identified and validated using an integrated analysis of bulk and single-cell RNA sequencing. Front Immunol 16, 1686948 (2025).

72. Dong, F., et al. ALKBH5 Facilitates Hypoxia-Induced Paraspeckle Assembly and IL8 Secretion to Generate an Immunosuppressive Tumor Microenvironment. Cancer Res 81, 5876–5888 (2021).

73. Du, W., et al. Histone lactylation-driven YTHDC1 promotes hepatocellular carcinoma progression via lipid metabolism remodeling. Cancer Lett 611, 217426 (2024).

74. Jia, Y., et al. Long non-coding RNA NEAT1 mediated RPRD1B stability facilitates fatty acid metabolism and lymph node metastasis via c-Jun/c-Fos/SREBP1 axis in gastric cancer. J Exp Clin Cancer Res 41, 287 (2022).

75. Krusnauskas, R., Stakaitis, R., Steponaitis, G., Almstrup, K. & Vaitkiene, P. Identification and comparison of m6A modifications in glioblastoma non-coding RNAs with MeRIP-seq and Nanopore dRNA-seq. Epigenetics 18, 2163365 (2023).

76. Lee, Q., et al. Overexpression of VIRMA confers vulnerability to breast cancers via the m(6)A-dependent regulation of unfolded protein response. Cell Mol Life Sci 80, 157 (2023).

77. Li, R., et al. The prognostic value and immune landscaps of m6A/m5C-related lncRNAs signature in the low grade glioma. BMC Bioinformatics 24, 274 (2023).

78. Liu, D., Ding, B., Liu, G. & Yang, Z. FUS and METTL3 collaborate to regulate RNA maturation, preventing unfolded protein response and promoting gastric cancer progression. Clin Exp Med 25, 15 (2024).

79. Liu, T., et al. Methyltransferase-like 14 suppresses growth and metastasis of renal cell carcinoma by decreasing long noncoding RNA NEAT1. Cancer Sci 113, 446–458 (2022).

80. Liu, Y., et al. The long non-coding RNA NEAT1 promotes the progression of human ovarian cancer through targeting miR-214-3p and regulating angiogenesis. J Ovarian Res 16, 219 (2023).

81. Mamontova, V., et al. NEAT1 promotes genome stability via m(6)A methylation-dependent regulation of CHD4. Genes & development 38, 915–930 (2024).

82. Peng, K., et al. ALKBH5 promotes the progression of infantile hemangioma through regulating the NEAT1/miR-378b/FOSL1 axis. Mol Cell Biochem 477, 1527–1540 (2022).

83. Qi, L., Yin, Y. & Sun, M. m6A-mediated lncRNA NEAT1 plays an oncogenic role in non-small cell lung cancer by upregulating the HMGA1 expression through binding miR-361-3p. Genes Genomics 45, 1537–1547 (2023).

84. Qin, X., et al. The disordered C terminus of ALKBH5 promotes phase separation and paraspeckles assembly. J Biol Chem 299, 105071 (2023).

85. Wang, J., et al. N6-methyladenosine reader hnRNPA2B1 recognizes and stabilizes NEAT1 to confer chemoresistance in gastric cancer. Cancer Commun (Lond) 44, 469–490 (2024).

86. Xu, Y., et al. LncRNA NEAT1 promotes immunosuppression in gastric cancer under endoplasmic reticulum stress by maintaining the M(6)A methylation of SEMA3A in CAFs. Mol Cancer (2026).

87. Yeermaike, A., Gu, P., Liu, D. & Nadire, T. LncRNA NEAT1 sponges miR-214 to promoted tumor growth in hepatocellular carcinoma. Mamm Genome 33, 525–533 (2022).

88. Zhang, J., et al. ALKBH5 promotes invasion and metastasis of gastric cancer by decreasing methylation of the lncRNA NEAT1. J Physiol Biochem 75, 379–389 (2019).

89. Zhang, S., Bai, Y., Wang, B. & Ju, H. ALKBH5 Accelerates the Progression of Head and Neck Squamous Cell Carcinoma by Decreasing Methylation of the lncRNA NEAT1. Appl Biochem Biotechnol 198, 1635–1655 (2026).

90. Zhou, Y., Zeng, P., Li, Y.H., Zhang, Z. & Cui, Q. SRAMP: prediction of mammalian N6-methyladenosine (m6A) sites based on sequence-derived features. Nucleic Acids Res 44, e91 (2016).

91. Qu, T., et al. Changes and relationship of N(6)-methyladenosine modification and long non-coding RNAs in oxidative damage induced by cadmium in pancreatic beta-cells. Toxicol Lett 343, 56–66 (2021).

92. Shu, B., Zhang, R.Z., Zhou, Y.X., He, C. & Yang, X. METTL3-mediated macrophage exosomal NEAT1 contributes to hepatic fibrosis progression through Sp1/TGF-beta1/Smad signaling pathway. Cell death discovery 8, 266 (2022).

93. Ma, D.B., Zhang, H., Wang, X.L. & Wu, Q.G. METTL3 aggravates cell damage induced by Streptococcus pneumoniae via the NEAT1/CTCF/MUC19 axis. Kaohsiung J Med Sci 40, 722–731 (2024).

94. Du, J., et al. Mesenchymal stem cell exosomes regulate TGFbeta/Smad3 by decreasing the METTL3-NEAT1 axis to inhibit scar progression after breast surgery. J Mol Histol 56, 161 (2025).

95. Chen, T., et al. Identification of long noncoding RNA NEAT1 as a key gene involved in the extramedullary disease of multiple myeloma by bioinformatics analysis. Hematology 28, 2164449 (2023).

96. Yankova, E., et al. Small-molecule inhibition of METTL3 as a strategy against myeloid leukaemia. Nature 593, 597–601 (2021).

97. Dong, P., et al. Long Non-coding RNA NEAT1: A Novel Target for Diagnosis and Therapy in Human Tumors. Frontiers in genetics 9, 471 (2018).

98. Yamazaki, T., et al. Functional Domains of NEAT1 Architectural lncRNA Induce Paraspeckle Assembly through Phase Separation. Molecular cell 70, 1038–1053 e1037 (2018).

99. Wu, Y. & Wang, H. LncRNA NEAT1 promotes dexamethasone resistance in multiple myeloma by targeting miR-193a/MCL1 pathway. J Biochem Mol Toxicol 32(2018).

100. Geng, W., et al. Resveratrol inhibits proliferation, migration and invasion of multiple myeloma cells via NEAT1-mediated Wnt/beta-catenin signaling pathway. Biomed Pharmacother 107, 484–494 (2018).

101. Yu, H., Peng, S., Chen, X., Han, S. & Luo, J. Long non-coding RNA NEAT1 serves as a novel biomarker for treatment response and survival profiles via microRNA-125a in multiple myeloma. J Clin Lab Anal 34, e23399 (2020).

102. Xu, H., Li, J. & Zhou, Z.G. NEAT1 promotes cell proliferation in multiple myeloma by activating PI3K/AKT pathway. European review for medical and pharmacological sciences 22, 6403–6411 (2018).

103. Taiana, E., et al. Long non-coding RNA NEAT1 shows high expression unrelated to molecular features and clinical outcome in multiple myeloma. Haematologica 104, e72–e76 (2019).

104. Taiana, E., et al. Long non-coding RNA NEAT1 targeting impairs the DNA repair machinery and triggers anti-tumor activity in multiple myeloma. Leukemia 34, 234–244 (2020).

105. Ghafouri-Fard, S. & Taheri, M. Nuclear Enriched Abundant Transcript 1 (NEAT1): A long non-coding RNA with diverse functions in tumorigenesis. Biomed Pharmacother 111, 51–59 (2019).

106. Gao, Y., et al. LncRNA NEAT1 sponges miR-214 to regulate M2 macrophage polarization by regulation of B7-H3 in multiple myeloma. Mol Immunol 117, 20–28 (2020).

107. Che, F., Ye, X., Wang, Y., Ma, S. & Wang, X. Lnc NEAT1/miR-29b-3p/Sp1 form a positive feedback loop and modulate bortezomib resistance in human multiple myeloma cells. Eur J Pharmacol 891, 173752 (2021).

108. Qin, J., et al. Aberrantly expressed long noncoding RNAs as potential prognostic biomarkers in newly diagnosed multiple myeloma: A systemic review and meta-analysis. Cancer Med 12, 2199–2218 (2023).

109. Wang, Q.M., et al. Exosomal lncRNA NEAT1 Inhibits NK-Cell Activity to Promote Multiple Myeloma Cell Immune Escape via an EZH2/PBX1 Axis. Mol Cancer Res 22, 125–136 (2024).

110. Wang, F., et al. LncRNA NEAT1 contributes to multiple myeloma progression via upregulating ARPC5 by sponging miR-133a. Arch Med Sci 21, 2088–2101 (2025).

111. Ren, Y., et al. Expression of NEAT1 can be used as a predictor for Dex resistance in multiple myeloma patients. BMC cancer 23, 630 (2023).

112. Puccio, N., et al. Combinatorial strategies targeting NEAT1 and AURKA as new potential therapeutic options for multiple myeloma. Haematologica 109, 4040–4055 (2024).

113. Xu, Y., et al. Long non-coding RNA NEAT1 promotes multiple myeloma malignant transformation via targeting miR-485-5p/ABCB8. Hematology 29, 2422153 (2024).

114. Wang, T., et al. LncRNA NEAT1 modulates myeloma cell autophagy and apoptosis by competitively binding miR-195-5p to regulate CEBPA. Discov Oncol 16, 2012 (2025).

115. Benini, G., et al. NEAT1: a multifaceted long non-coding RNA in multiple myeloma. Haematologica (2025).

116. Dierks, D. & Schwartz, S. Why m(6)A? An RNA surveillance model. Molecular cell (2026).

